# How important are functional and developmental constraints on phenotypic evolution? An empirical test with the stomatal anatomy of flowering plants

**DOI:** 10.1101/2021.09.02.457988

**Authors:** Christopher D. Muir, Miquel Àngel Conesa, Jeroni Galmés, Varsha S. Pathare, Patricia Rivera, Rosana López Rodríguez, Teresa Terrazas, Dongliang Xiong

## Abstract

Functional and developmental constraints on phenotypic variation may cause traits to covary over millions of years and slow populations from reaching their adaptive optima. Alternatively, trait covariation may result from selective constraint if some trait combinations are generally maladaptive. Quantifying the relative contribution of functional, developmental, and selective constraints on phenotypic variation is a longstanding goal of macroevolution, but it is often difficult to distinguish different types of constraints. The anatomy of leaves with stomata on both surfaces (amphistomatous) present a unique opportunity to test the importance of functional and developmental constraints on phenotypyic evolution. The key insight is that stomata on each leaf surface encounter the same functional and developmental constraints, but potentially different selective constraints because of leaf asymmetry in light capture, gas exchange, and other features. Independent evolution of stomatal traits on each surface imply that functional and developmental constraints alone likely do not explain trait covariance. Packing limits on how many stomata can fit into a finite epidermis and cell-size-mediated developmental integration are hypothesized to constrain variation in stomatal anatomy. The simple geometry of the planar leaf surface and knowledge of stomatal development make it possible to derive equations for phenotypic (co)variance caused by these constraints and compare them with data. We analyzed evolutionary covariance between stomatal density and length in amphistomatous leaves from 236 phylogenetically independent contrasts using a robust Bayesian model. Stomatal anatomy on each surface diverges partially independently, meaning that packing limits and developmental integration are not sufficient to explain phenotypic (co)variation. Hence, selective constraints, which require an adaptive explanation, likely contribute to (co)variation in ecologically important traits like stomata. We show how it is possible to evaluate the contribution of different constraints by deriving expected patterns of (co)variance and testing them using similar but separate tissues, organs, or sexes.

## Introduction

The ability for traits to evolve independently of one another is a necessary prerequisite for populations to ascend a fitness peak in multivariate trait space (Lewontin 1978). If traits can evolve independently and there is sufficient genetic variation, then selection should move populations toward their multivariate phenotypic optimum. Yet divergence in one trait often covaries with other traits and covariance can persist for millions of years (Schluter 1996). Covariance is “a rough local measure of the strength of constraint” (Maynard Smith et al. 1985) that can be broken down into functional, developmental, and selective constraints. Functional constraints are “limitations imposed by time, energy, or the laws of physics” (Arnold 1992). In other words, certain trait combinations are not physically or geometrically possible. A classic example is shell coiling among invertebrate lineages in which the morphospace of possible phenotypes is constrained by hard geometrical limits (Raup 1966; McGhee 1999). Within the space of possible phenotypes, developmental constraints can “bias... the production of variant phenotypes or [place] a limitation on phenotypic variability” (Maynard Smith et al. 1985). For example, Fibonacci phyllotaxis may arise from a packing constraint of primordia on the developing apex (Mitchison 1977; Reinhardt and Gola 2022; but see Niklas 1988 for an adaptive explanation). Natural selection constrains phenotypic variation by preventing maladaptive forms from evolving. Selection causes trait covariation because ‘missing’ trait combinations are maladaptive. Understanding phenotypic constraint is challenging, but a useful starting point is determining whether phenotypic covariation can be explained by functional or developmental constraints (McGhee 1999, 2007; Olson 2019). If phenotypic covariation is inconsistent with functional and developmental constraints, this provides a strong impetus to test for selective constraint. Selective constraint does not mean that populations do not respond to selection, but rather that limits on phenotypic variation arise because selection never favors certain trait values.

In this study, we will address packing problems and developmental integration, specific forms of functional and developmental constraint relevant to our study system, stomatal anatomy. We introduce these concepts generally in this paragraph. Packing a number objects into a finite space is a common functional constraint on organisms. Regular geometries that appear in nature such as helices and hexagons (think DNA and honeycombs) are often optimal solutions to packing problems (Mackenzie 1999; Maritan et al. 2000). Notice that functional constraint does not preclude selection, but the presence of a packing limit changes the range of possible phenotypes. Developmental integration is a form of developmental constraint on multivariate phenotypic evolution and we use these terms interchangeably in this study. Developmentally integrated traits have a “disposition for covariation” (Armbruster et al. 2014), meaning that evolutionary divergence between lineages in one trait will be tightly associated with divergence in another trait. Allometry is a classic, albeit contested, example of developmental integration that may constrain phenotypic evolution (reviewed in Pélabon et al. 2014). Strong allometric covariation between traits within populations can constrain macroevolutionary divergence for long periods of time depending on the strength and direction of selection (Lande 1979). However, developmental integration does not necessarily hamper adaptation, and can even accelerate adaptive evolution when trait covariation is aligned with the direction of selection (Hansen 2003). For example, fusion of floral parts increases their developmental integration which may increase the rate and precision of multivariate adaptation to specialist pollinators (Berg’s rule, Berg 1959, 1960; Conner and Lande 2014; Armbruster et al. 1999).

Biologists have studied phenotypic constraints for decades, but progress is challenging because many multivariate phenotypes are too complex or too poorly understood to quantitatively distinguish functional, developmental, and selective constraints over macroevolutionary timescales. Stomatal anatomy on the leaves of flowering plants provide an exceptional opportunity because 1) there are 1000s of species to compare and 2) the main packing constraints and developmental steps are analytically tractable. This means it is possible to derive quantitative predictions and test their generality using large comparative data sets representing millions of years of evolutionary history. For this purpose, a heretofore unappreciated fact about stomata is that many leaves have stomata on both lower and upper surfaces. The packing and developmental constraints are the same for stomata on each surface, but the selective constraints may differ. Therefore, if packing and developmental constraints dominate, stomatal anatomy on each surface should diverge in concert. Failure to do so implies that independent evolution is possible and that selective constraints most likely explain at least some of the covariance between traits. Phylogenetic comparisons of stomatal anatomy provide a statistically powerful, general, and elegant way to distinguish different phenotypic constraints that would be impossible in many other traits with as much ecological significance. The next sections provide background information on stomatal anatomy, how it varies, and why functional or developmental constraints might be important.

### The adaptive significance and ecological distribution of variation in stomatal anatomy

Stomata are microscopic pores formed by a pair of guard cells that regulate gas exchange (CO_2_ gain and water vapor loss) on the leaves or other photosynthetic surfaces of most land plants. Stomata originated once in the history of land plants around 500 Ma, diversified rapidly in density and size, and have been maintained in most lineages except some bryophytes and aquatic plants (recently reviewed in Clark et al. 2022). Stomata respond physiologically by opening and closing in response to light, humidity, temperature, circadian rhythm, and plant water status (Hetherington and Woodward 2003; Lawson and Matthews 2020). The stomatal size, density, and distribution on a mature leaf do not change, so the maximum rate of gas exchange is fixed. However, the plant may respond plastically to environmental cues such as light and CO_2_ by altering stomatal anatomy in new leaves (Casson and Gray 2008). Physiological responses (aperture change) and plastic responses (new leaves with changed anatomy) may be alternative strategies for plants to acclimate to environmental change (Haworth, Elliott-Kingston, and McElwain 2013). Finally, stomatal anatomy can evolve due to inherited changes in stomatal development. Plastic and genetic changes in stomatal anatomy are both ecologically important, but most studies do not use a common garden design that would tease apart their relative contribution.

We focus on anatomical variation in the density, size, and patterning of stomata on a leaf because these factors set the maximum stomatal conductance to CO_2_ diffusing into a leaf and the amount of water that transpires from it (Sack et al. 2003; Franks and Farquhar 2001; Galmés et al. 2013; Harrison et al. 2020). Plants typically operate below their anatomical maximum by dynamically regulating stomatal aperture. Even though operational stomatal conductance determines the realized photosynthetic rate and water-use efficiency, anatomical parameters are useful in that they set the range of stomatal function (de Boer et al. 2016) and are correlated with actual stomatal function under natural conditions (Murray et al. 2020). All else being equal, larger, more densely packed, but evenly spaced stomata increase gas exchange (Franks and Beerling 2009; Dow, Berry, and Bergmann 2014; Lehmann and Or 2015). Smaller stomata may also be able to respond more rapidly than larger stomata, proving the ability of leaves to track short duration environmental change (Drake, Froend, and Franks 2013). Stomata are most often found only on the lower leaf surface (hypostomy), but occur on both surfaces (amphistomy) in some species (Metcalfe and Chalk 1950; Parkhurst 1978; Mott, Gibson, and O’Leary 1982). Amphistomatous leaves have a second parallel pathway from the substomatal cavities through the leaf internal airspace to sites of carboxylation in the mesophyll (Parkhurst 1978; Gutschick 1984). Thus amphistomatous leaves have lower resistance to diffusion through the airspace which increases the photosynthetic rate (Parkhurst and Mott 1990). If total stomatal and other conductances to CO_2_ supply could be held constant, then an amphistomatous leaf will have a greater conductance than an otherwise identical hypostomatous leaf. The magnitude of the advantage depends on the resistance to diffusion through the internal airspace, which is variable among species.

The adaptive significance and ecological distribution of leaves with different stomatal anatomies is complex and there is much yet to learn. Seed plants posses a wider range of stomatal anatomies than ferns and fern allies, which are restricted to having large stomata, at low density, only on the lower surface (de Boer et al. 2016). In general, trees and shrubs have greater stomatal density than herbs, but there is a of lot variation within growth forms depending on the ecological niche (Salisbury 1928; Kelly and Beerling 1995). A commonly observed trend is that leaves from higher light environments tend to have greater stomatal density (Salisbury 1928; Mott, Gibson, and O’Leary 1982; Gibson 1996; W. K. Smith, Bell, and Shepherd 1998; Jordan, Carpenter, and Brodribb 2014; Muir 2015; Bucher et al. 2017). This may explain why, perhaps counterintuitively, plants in dry environments tend to have more stomata. Drier habitats are more open, enabling plants with higher stomatal density to photosynthesize more when water is available, but close stomata during drought (Liu et al. 2018). Over recent human history, stomatal density has tended to decline within species as atmospheric CO_2_ concentrations have risen (Woodward 1987; Royer 2001). It is unclear whether most of this change is plastic or genetic, but the overall direction is consistent with the hypothesis that plants decrease gas exchange as CO_2_ availability increases.

Stomatal ratio is a continuous measure of how stomata are deployed across leaf surfaces. The adaptive significance of variation in stomatal ratio is uncertain, but we have some clues based on the distribution of hypo- and amphistomatous leaves. Despite the fact that amphistomy can increase photosynthesis, most leaves are hypostomatous. Amphistomy should increase photosynthesis most under saturating-light conditions where CO_2_ supply limits photosynthesis. However, the light environment alone cannot explain why hypostomatous leaves predominate in shade plants (Muir 2019), suggesting that we need to understand the costs of upper stomata better. One factor may be increased vulnerability to pathogens. For example, upper stomata increase susceptibility to rust pathogens in *Populus* (McKown et al. 2014, 2019; Fetter, Nelson, and Keller 2021). Amphistomy may also cause the palisade mesophyll to dry out under strong vapor pressure deficits (Buckley et al. 2015). Other hypotheses about the adaptive significance of stomatal ratio are discussed in Muir (2015) and Drake et al. (2019).

### Major features of stomatal anatomical macroevolution

Two major features of stomatal anatomy have been recognized for decades but we do not yet understand the evolutionary forces that generate and maintain them. We denote these two features as “inverse size-density scaling” and “bimodal stomatal ratio” (Fig. 1). Inverse size-density scaling refers to the negative interspecific correlation between the size of the stomatal apparatus and the density of stomata (Weiss 1865; Franks and Beerling 2009; de Boer et al. 2016; Sack and Buckley 2016; Liu et al. 2021). Across species, leaves with smaller stomata tend to pack them more densely, but there is significant variation about this general trend (Fig. 1a). Stomatal size and density determine the maximum stomatal conductance to CO_2_ and water vapor but also take up space on the epidermis, which could be costly for both construction and maintenance. Natural selection should favor leaves that have enough stomata of sufficient size to supply CO_2_ for photosynthesis. Hence leaves with few, small stomata and high photosynthetic rates do not exist because they would not supply enough CO_2_. Conversely, excess stomata or extra large stomata beyond the optimum may result in stomatal interference where the CO_2_ concentration gradient around one stomate merges with that of its neighbor (Zeiger, Farquhar, and Cowan 1987; Lehmann and Or 2015), incur metabolic costs (Deans et al. 2020), and/or risk hydraulic failure (Henry et al. 2019). The distribution of stomatal size and density may therefore represent the combinations that ensure enough, but not too much, stomatal conductance. Franks and Beerling (2009) further hypothesized that the evolution of small stomata in angiosperms enabled increased stomatal conductance while minimizing the epidermal area allocated to stomata.

**Figure 1:**
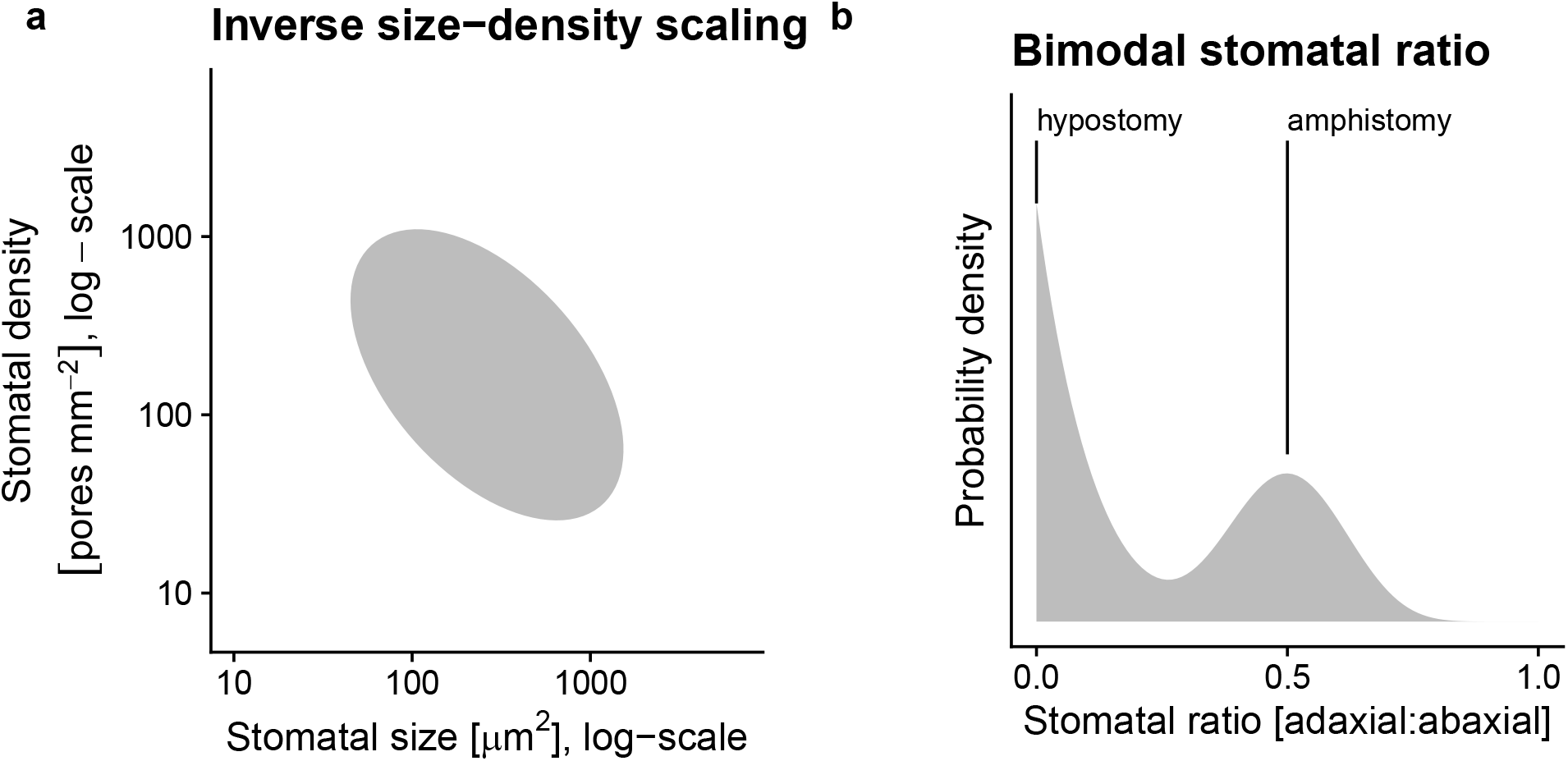
Two salient features of stomatal anatomy in flowering plants are the (a) inverse relationship between stomatal size and density and (b) the bimodal distribution of stomatal ratio. At broad phylogenetic scales, species with smaller stomata on their leaves (x-axis, log-scale) tend to have greater stomatal density (x-axis, log-scale), but there is a lot of variation about the overall trend indicated by the grey ellipse. Hypostomatous leaves (stomatal ratio = 0) are more common than amphistomatoues leaves, but within amphistomatous leaves, the density of stomata on each surface tends to be similar (stomatal ratio ≈ 0.5), which we refer to as bimodal stomatal ratio.

A striking feature of the interspecific variation in stomatal ratio is that trait values are not uniformly distributed, but strongly bimodal (Fig. 1b). Bimodal stomatal ratio refers to the observation that the ratio of stomatal density on the adaxial (upper) surface to the density on the abaxial (lower) has distinct modes (Fig. 1b). Amphistomy occurs most often in herbaceous plants from open, high light habitats (Salisbury 1928; Mott, Gibson, and O’Leary 1982; Gibson 1996; W. K. Smith, Bell, and Shepherd 1998; Jordan, Carpenter, and Brodribb 2014; Muir 2015, 2018; Bucher et al. 2017). Muir (2015) described bimodal stomatal ratio formally but the pattern is apparent in earlier comparative studies of the British flora (cf.Peat and Fitter 1994, fig. 1).

### Packing limits, developmental integration, and stomatal anatomy

Given the significance of stomata for plant function and global vegetation modeling (Berry, Beerling, and Franks 2010), we would like to understand what factors constrain their anatomical variation. Here we focus on interspecific variation in mean trait values rather than intraspecifc variation. Packing constraints and developmental integration could explain inverse size-density scaling. The density and size of a stomata on a planar leaf surface can be viewed as a packing problem where the total area allocated to stomata cannot exceed the total leaf area. This is a functional, or geometric, constraint because certain combinations of large size and high density are not physically possible. This functional constraint cannot explain why combinations of low density and small size are rare, but may explain why stomatal size must decrease when density increases as the leaf runs out of space. The packing limit of functional stomata is less than the entire leaf area, but the exact value is unclear. The realized upper limit is close to 1/3 or 1/2 (de Boer et al. 2016; Sack and Buckley 2016; Liu et al. 2021) for the species’ mean, not an individual leaf.

Guard cell size and spacing between stomata (the inverse of density) are developmentally intertwined because guard cells and epidermal pavement cells between stomata develop from the same meristem. Before guard cell meristemoids form via asymmetric cell division (Dow and Bergmann 2014), the size of guard and epidermal cells are influenced by meristematic cell volume and expansion. Evolutionary shifts in meristematic cell volume or expansion rate could cause both increased stomatal size and lower density because epidermal cells between stomata are larger (Brodribb, Jordan, and Carpenter 2013). For example, larger genomes increase meristematic cell volume (Šímová and Herben 2012), which sets a lower bound on final cell volume. Although different expansion rates in guard and epidermal pavement cells can reduce the correlation in their final size, the fact that species with larger genomes tend toward having larger stomata and lower density may indicate an effect of development integration on stomatal anatomy (Beaulieu et al. 2008; Simonin and Roddy 2018; Roddy et al. 2020). Developmental integration in this case would not necessarily hinder adaptive evolution if the main axes of selection were aligned with the developmental correlation. For example, if higher maximum stomatal conductance were achieved primarily by increasing stomatal density and decreasing stomatal size as proposed by Franks and Beerling (2009), then developmental integration might accelerate the response to selection compared to a case where stomatal size and density are completely independent.

Muir (2015) derived general conditions in which bimodality arises because adaptive optima are restricted to separate regimes, but this model has not been tested. An alternative hypothesis is that stomatal traits on the ab- and adaxial surfaces are developmentally integrated because stomatal development is regulated the same way on each surface. In hypostomatous leaves, stomatal development is turned off in the adaxial surface. In amphistomatous leaves, stomatal development proceeds on both surfaces, but evolutionary changes in stomatal development affect traits on both surfaces because they are tethered by a shared developmental program. This is a developmental constraint because the fact that stomatal development is the same on each surface constrains the type of variation available for selection. Developmental integration would lead to a bimodal trait distribution because leaves would either be hypostomatous (stomatal ratio equal to 0) or have similar densities on each surface (stomatal ratio approximately 0.5). To our knowledge, this hypothesis has not been put forward in the literature.

### Hypotheses and predictions

The overarching question is whether major features of stomatal anatomy in terrestrial angiosperms are consistent with packing constraints and/or developmental integration mediated by cell size. Since stomata on both surfaces of amphistomatous leaves are subject to the same functional and developmental constraints, if these constraints are most important we predict similar patterns of trait covariation on abaxial and adaxial surfaces. Conversely, if traits covary differently on each surface it would indicate that stomatal anatomical traits can evolve independently and selective constraints likely contribute to covariation. Analogously, variation in the genetic correlation and interspecific divergence of sexually dimorphic traits in dioecious species demonstrate that integration is not fixed and can be modified by selection (Barrett and Hough 2013). We framed specific hypotheses and predictions around how functional or developmental constraints might explain either inverse size-density scaling or bimodal stomatal ratio.

#### Inverse size-density scaling

Both packing limits and developmental integration could contribute to inverse size-density scaling. If limits on the fraction of epidermal area occupied by stomata constrains the combinations of stomatal size and density that are evolutionarily accessible, then we predict that evolutionary divergence in stomatal size and/or density will decrease as the fraction of epidermal area occupied by stomata increases. Furthermore, if divergence slows as epidermal area occupied by stomata because of a packing limit, it should slow down the same way for both ab- and adaxial surfaces.

The second hypothesis is the cell size mediates developmental integration between stomatal size and density. If developmental integration is the primary reason for inverse size-density scaling, then amphistomatous leaves will exhibit identical size-density scaling on each surface. If the stomatal size and density scale differently on each surface, this implies that they can evolve independently and that selective constraints likely explain some of their covariance. Furthermore, we predicted that divergence in genome size, which is strongly associated with meristematic cell volume (Šímová and Herben 2012), would covary with stomatal size and density similarly on each surface.

#### Bimodal stomatal ratio

If the developmental integration hypothesis is correct, it also implies stomatal size and density will diverge in concert on each surface because the developmental function is fixed. Therefore we predict that divergence of stomatal traits on one surface will be isometric with divergence in stomatal traits on the other surface. This type of developmental integration limits the expression of variation and could give rise to a bimodal stomatal ratio. Suppose that in hypostomatous leaves, stomatal development is completely suppressed. In amphistomatous leaves, stomatal development proceeds identically on each surface because the developmental function is identical. This would lead to a tendency for equal density on each surface.

We formalized these hypotheses into a mathematical framework to derive quantitative predictions that we tested in a phylogenetic comparative framework by compiling stomatal anatomy data from the literature for a broad range of flowering plants.

## Materials and Methods

Unless otherwise mentioned, we performed all data wrangling and statistical analyses in *R* version 4.2.1 (R Core Team 2022). Source code is publicly available on GitHub (https://github.com/cdmuir/stomata-independence) and will be archived on Zenodo upon publication.

### Theory: divergence with and without developmental constraint

Developmental integration could shape patterns of phenotypic macroevolution, but a major hindrance to progress is that verbal models do not make precise, quantitative predictions that distinguish it from alternatives. An advantage of testing developmental integration in stomata is that their development is well studied (Bergmann and Sack 2007; Dow and Bergmann 2014; Sack and Buckley 2016). We can leverage that knowledge to build a developmental function and derive equations for phenotypic (co)variance caused by developmental integration. If observed patterns of evolution are inconsistent with developmental integration, theory may also help identify which parameters lead to developmental disintegration. We combined and extended stomatal development models to predict how stomatal density and length would diverge if stomatal development were constrained and how those predictions would change if stomatal development were unconstrained. We summarize our methods verbally here and direct readers interested in the mathematical details to Notes A1. A graphical summary is provided in Fig. A3. We imposed constraint by assuming the stomatal developmental function is constrained. The developmental function maps cell size prior to differentiation onto stomatal size and density using two parameters. The first parameter describes how cell volume is apportioned between epidermal cells and guard cell meristemoids during asymmetric cell division (Dow and Bergmann 2014). The second parameter is stomatal index (Salisbury 1928; Sack and Buckley 2016), which is determined by amplifying and spacing divisions after asymmetric cell division (Dow and Bergmann 2014). When these parameters are fixed, divergence in stomatal size and density is determined by divergence in meristematic cell volume and expansion prior to asymmetric division. We relaxed this constraint by treating parameters of the developmental function as random variables that can diverge between species. We used random variable algebra to derive predicted (co)variance in divergence between stomatal length and density (see Lynch and Walsh 1998 for key random variable algebra theorems).

### Data synthesis

We searched the literature for studies that measured stomatal density and stomatal size, either guard cell length or stomatal pore length, for both abaxial and adaxial leaf surfaces. In other words, we did not include studies unless they reported separate density and size values for each surface. We did not record leaf angle because it is typically not reported, but we presume that for the vast majority of taxa that the abaxial is the lower surface and the adaxial is the upper surface. This is reversed in resupinate leaves, but to the best of our knowledge, our synthesis did not include resupinate leaves. None of the species with resupinate leaves listed by Chitwood et al. (2012) are in our data set. We refer to guard cell length as stomatal length and converted stomatal pore length to stomatal length assuming guard cell length is twice pore length (Sack and Buckley 2016). Table 1 lists focal traits and symbols.

**Table 1:**
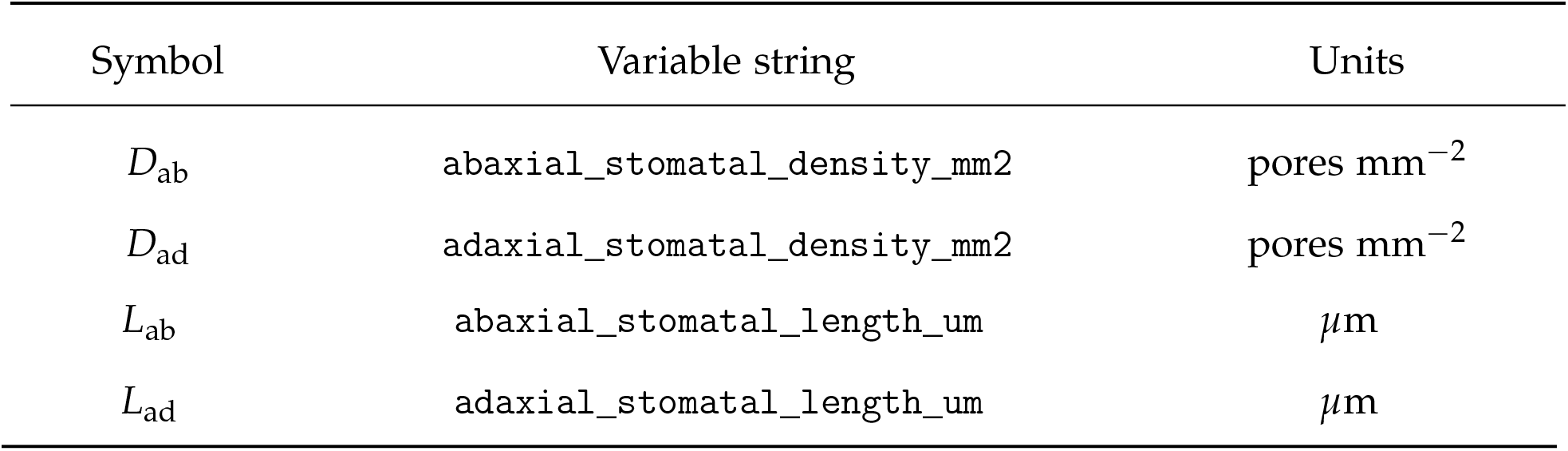
Stomatal anatomical traits with mathemtical symbol, variable string used in source code, and scientific units.

| Symbol | Variable string | Units |
| --- | --- | --- |
| $D_{ab}$ | abaxial_stomatal_density_mm2 | pores $\text{mm}^{-2}$ |
| $D_{ad}$ | adaxial_stomatal_density_mm2 | pores $\text{mm}^{-2}$ |
| $L_{ab}$ | abaxial_stomatal_length_um | $\mu\text{m}$ |
| $L_{ad}$ | adaxial_stomatal_length_um | $\mu\text{m}$ |

Data on stomatal anatomy are spread over a disparate literature and we have not attempted an exhaustive synthesis of amphistomatous leaf stomatal anatomy. We began our search by reviewing papers that cited key studies of amphistomy (Parkhurst 1978; Mott, Gibson, and O’Leary 1982; Muir 2015). We supplemented these by searching Clarivate *Web of Science* for “guard cell length” because most studies that report guard cell length also report stomatal density, whereas the reverse is not true. We identified additional studies by reviewing the literature cited of papers we found and through opportunistic discovery. The final data set contained 5104 observations of stomatal density and length from 1242 taxa and 38 primary studies (Table A5). However, many of these data were excluded if taxonomic name and phylogenetic placement could not be resolved (see below). Finally, we included some unpublished data. Stomatal size data were collected on grass species described in Pathare, Koteyeva, and Cousins (2020). We also included unpublished data on 14 amphistomatous wild tomato species (*Solanum* sect. *Lycopersicum* and sect. *Lycopersicoides*) grown in pots under outdoor summer Mediterranean conditions as described in Muir, Galmés, and Conesa (2022). We took ab- and adaxial epidermal imprints using clear nail polish of the mid-portion of the lamina away from major veins on the terminal leaflet of the youngest, fully expanded leaf from 1-5 replicates per taxon. With a brightfield light microscope, we counted stomata in three 0.571 mm^2^ fields of view and divided by the total area to estimate density. We measured the average guard cell length of 60 stomata, 20 per field of view, to estimate stomatal size. The data set is publicly available as an *R* package **ropenstomata** (https://github.com/cdmuir/ropenstomata).

We collected data on genome size from the Angiosperm DNA C-values database (Leitch et al. 2019; Pellicer and Leitch 2020). When multiple ploidy levels were available for a taxon, we chose the lowest one for consistency.

### Phylogenetically independent contrasts

We generated an ultrametric, bifurcating phylogeny of 638 taxa by resolving and removing ambiguous taxonomic names, placing taxa on the GBOTB.extended mega-tree of seed plants (S. A. Smith and Brown 2018; Zanne et al. 2014), and resolving polytomies using published sequence data. Divergence times in this phylogeny are based on extensive fossil calibration (see S. A. Smith and Brown 2018; Magallón et al. 2015 for further detail). The complete methodology is described in the online supplement (Notes A2).

From this phylogeny, we extracted 236 phylogenetically independent taxon pairs (Table A1). A fully resolved, bifurcating four-taxon phylogeny can have two basic topologies: ((`A,B`), (`C,D`)) or ((`A,B`),`C`),`D`)). Taxon pairs include all comparisons of A with B and C with D in each four-taxon clade. We extracted pairs using the `extract_sisters()` function in R package **diverge** version 2.0.4 (Anderson and Weir 2021) and custom scripts (see source code). Taxon pairs are the most closely related pairs in our data set, but they are mostly not sister taxa in the sense of being the two most closely related taxa in the tree of life. For each pair we calculated phylogenetically independent contrasts (Felsenstein 1985) as the difference in the log_10_-transformed trait value (see Beaulieu et al. 2008 for a similar approach). Contrasts are denoted as Δlog(trait). We log-transformed traits for normality because like many morphological and anatomical traits they are strongly right-skewed. Log-transformation also helps compare density and length, which are measured on different scales, because log-transformed values quantify proportional rather than absolute divergence.

### Parameter estimation

All hypotheses make predictions about trait (co)variance matrices or parameters derived from them (see Notes A1 and subsections below). Within and among species covariation is a hallmark of developmental integration (Armbruster 1988), but other evolutionary processes also lead to covariance. Distinguishing between them requires deriving predictions and testing whether observed covariance is consistent with one hypothesis or another. We estimated the 4 × 4 covariance matrix of phylogenetically independent contrasts between log-transformed values of Δlog(*D*_ab_), Δlog(*D*_ad_), Δlog(*L*_ab_), and Δlog(*L*_ad_) using a distributional multiresponse robust Bayesian approach. See Table 1 for variable definitions. We denote variances as Var[Δlog(trait)] and covariances as Cov[Δlog(trait_1_), Δlog(trait_2_)]. We used a multivariate *t*-distribution rather than a Normal distribution because estimates using the former are more robust to exceptional trait values (Lange, Little, and Taylor 1989). We also estimated whether the variance in trait divergence increases with time. Under many trait evolution models (e.g. Brownian motion), interspecific variance increases through time. To account for this, we included time since taxon-pair divergence as an explanatory variable affecting the trait covariance matrix.

For the packing limit hypothesis, we tested whether the variance in stomatal trait divergence, Var[Δ log(trait)], decreases as stomatal allocation increases. The fraction of epidermal area allocated to stomata (*f_S_*) is the product of stomatal density and area occupied by a stomatal apparatus. Because guard cell shape is similar in most plant lineages except grasses, the area can be well approximated from guard cell length as *A* = *jL*^2^ where *j* = 0.5 for most species with kidney-shaped guard cells and *j* = 0.125 for grasses with dumbbell-shaped guard cells (Sack and Buckley 2016). For each contrast, we calculated the average *f_S_* on each surface between those two taxa for use as our explanatory variable. The statistical model allowed the effect of *f_S_* on Var[Δ log(trait)] to vary between traits and leaf surfaces. We also included time since taxon-pair divergence as an explanatory variable and used a multivariate *t*-distribution as described above.

We fit all models in Stan 2.29 (Stan Development Team 2022) using the R packages **brms** version 2.17.0 (Bürkner 2017, 2018) with a **cmdstanr** version 0.5.2 backend (Gabry and Češnovar 2022). It ran on 2 parallel chains for 1000 warm-up iterations and 1000 sampling iterations. All parameters converged 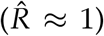 and the effective sample size from the posterior exceeded 1000 (Vehtari et al. 2021). We used the posterior median for point estimates and calculated uncertainty with the 95% highest posterior density (HPD) interval from the posterior distribution.

### Hypothesis testing

#### Does divergence slow as epidermal space fills?

We tested the packing limit hypothesis by estimating the effect of *f_S_* on Var[Δ log(trait)] for each trait and leaf surface. If there is an upper bound on *f_S_*, we predict the effect of *f_S_* on Var[Δ log(trait)] will be < 0. Specifically, the 95% HPD intervals should not include 0. Further, the coefficient should be the same on each surface, so the 95% HPD intervals for difference should encompass 0.

#### Is size-density scaling the same on both leaf surfaces?

We tested whether the covariance between divergence in stomatal length and stomatal density on each leaf surface is the same. If size and density are developmentally integrated, we predict the covariance matrices will not be significantly different. Specifically, the 95%

HPD intervals of the difference in covariance parameters should not include 0 if:

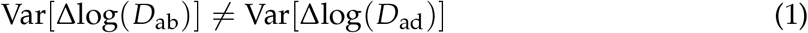

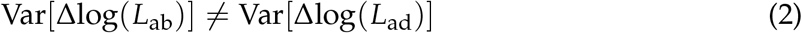

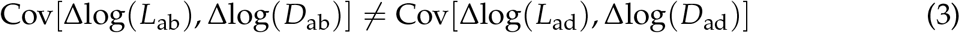

#### Do abaxial and adaxial stomatal traits evolve isometrically?

If stomatal traits on each surface are developmentally integrated then divergence in the trait on one surface should result in a 1:1 (isometric) change in the trait on the other surface. Furthermore, there should be relatively little variation away from a 1:1 relationship. Conversely, if traits can evolve independently then the change in the trait on one surface should be uncorrelated with changes on the other. We tested for isometry by estimating the standardized major axis (SMA) slope of divergence in the abaxial trait against divergence in the adaxial trait for both stomatal length and stomatal density. If change on each surface is isometric, then the HPD intervals for the slope should include 1. We used the coefficient of determination, *r*^2^, to quantify the strength of integration, where a value of 1 is complete integration and a value of 0 is complete disintegration.

## Results

### Theory: from developmental integration to disintegration

We asked how divergence in stomatal length and density would covary if the developmental function were constrained and compared it to their divergence when the developmental function can evolve. When the developmental function is constrained this means that allocation to guard cell meristemoids during asymmetric division and stomatal index are fixed (see Notes A1 for mathematical description). Under these assumptions, divergence in stomatal length and density is mediated entirely by divergence in meristematic cell volume and expansion prior to differentiation. Developmental integration is strong because divergence in density is perfectly negatively correlated with divergence in size. In contrast, stomatal length and density can diverge independently when the developmental function is not fixed. Divergence in asymmetric cell division affects stomatal size independently of density; divergence in stomatal index affects stomatal density independently of size. Divergence in the developmental function causes developmental disintegration because stomatal density and size can diverge independently. Developmental integration is minimal when asymmetric cell division and/or stomatal index diverge more than meristematic cell volume and expansion. The three main conclusions are that 1) developmental constraint leads to developmental integration; 2) different (co)variance in divergence of stomatal length and density on each surface implies the developmental function is not fixed; and 3) divergence in different components of the developmental function affect stomatal length and density differently. See Notes A1 and Table A4 for more a complete derivation and detailed predictions.

### Divergence in stomatal traits slows as fS increases

The variance in trait divergence decreases as the fraction of epidermal area allocated to stomata, *f_S_*, increases (Figs. 2, A1). The effect of *f_S_* was strongest for *D*_ad_ and 95% HPD intervals did not overlap 0 for 3 of 4 comparisons (Table A2). Variance in divergence for *D*_ad_ declined more rapidly with *f_S_* than that for *D*_ab_ (difference and 95% HPD interval in slope, log-link scale: −10.7 [-17.2,-4.2]). Variance in length divergence declined similarly with *f_S_* on both surfaces (difference and 95% HPD interval in slope, log-link scale: −2.8 [-9.6,4]).

**Figure 2:**
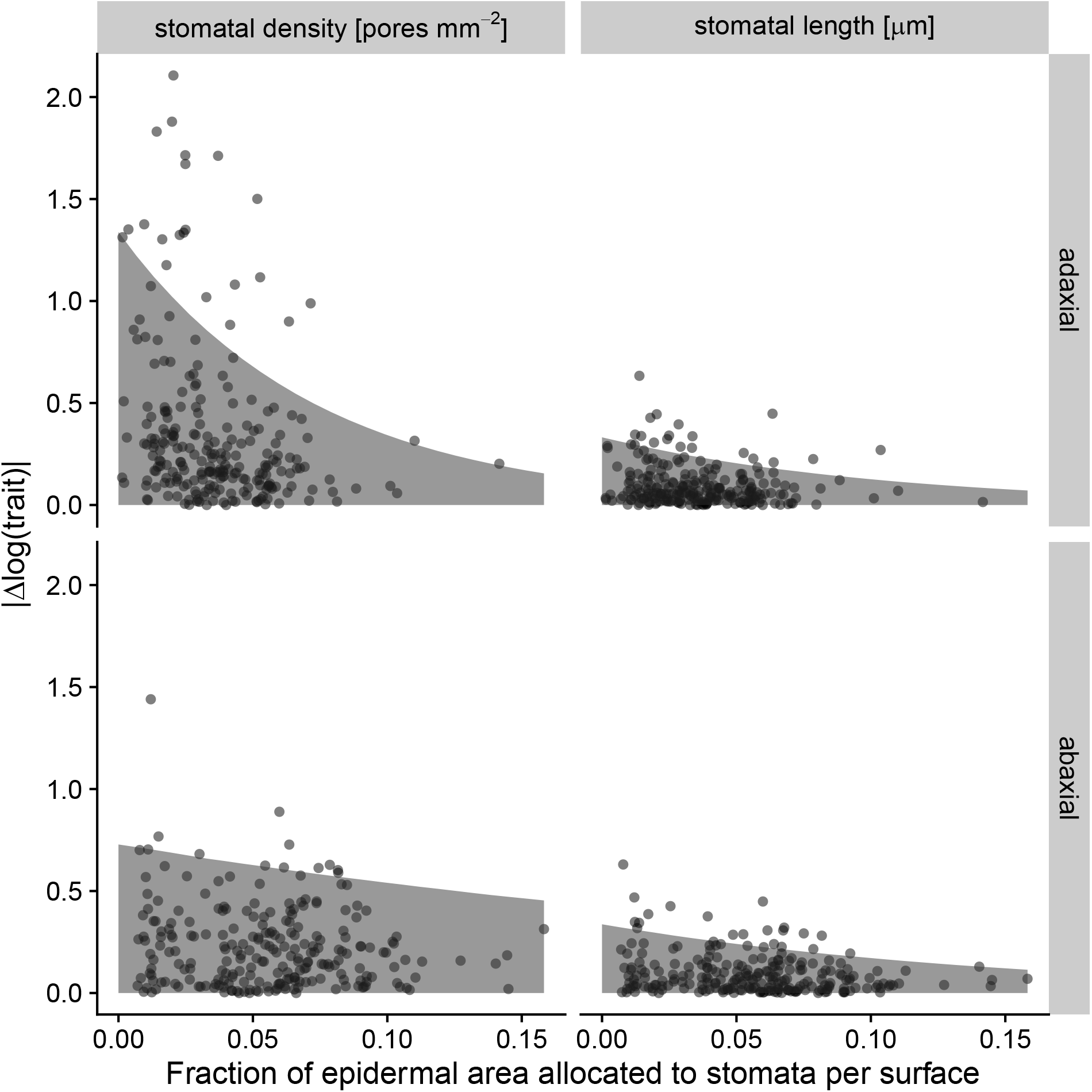
Evolutionary divergence slows down as epidermal space fills up. This pattern is consistent with functional constraints (packing limits) constraining evolution, but only when stomata occupy a large fraction of epidermal area. The shaded area in each facet indicates the estimate of where 95% of the 236 phylogenetically independent contrasts fall as a function of the fraction of epidermal area allocated to stomata per surface. Each point is the absolute value of Δ log(trait) for stomatal density (left facets) or length (right facets) on the adaxial (upper facets) and abaxial (lower facets) surface. The fraction of epidermal area allocated to stomata is the average value per surface between the two taxa in each contrast. Divergence in anatomical traits is more variable when stomata occupy a smaller area, especially for adaxial stomatal density (upper left facet). See Table A2 for all parameter estimates and confidence intervals.

### Adaxial stomatal density is more variable, but size-density covariance is similar on both surfaces

Stomatal length negatively covaries with stomatal density similarly on both surfaces, but on the adaxial surface there are many more taxa that have low stomatal density and small size compared to the abaxial surface (Fig. 3). In principle, this pattern could arise either because size-density covariance differs or the variance in adaxial stomatal density increases faster than that for abaxial stomatal density. The interspecific variance increases with time since divergence for all traits (Table A3). For consistency, we therefore report estimates conditional on time since divergence set to 0. Across pairs, we estimate that the covariance between size and density is similar. The median estimate is Cov[Δlog(*L*_ad_), Δlog(*D*_ad_)] – Cov[Δlog(),Δlog(*D*_ab_)] = 3.18 × 10^−4^, but 0 is within the range of uncertainty (95% HPD interval [–3.85 × 10^−3^,4.26 × 10^−3^]). However the variance in adaxial stomatal density is significantly greater than the abaxial stomatal density [Fig. 4]). We estimate Var[Δlog(*D*_ad_)] is 4.00 × 10^−2^ (95% HPD interval [1.42 × 10 ^−2^,7.07 × 10 ^−2^]) greater than Var[Δlog(*D*_ab_)]. The variance in stomatal length was similar for both surfaces, with an estimate of −4.54 × 10^-4^ (95% HPD interval [-1.67 × 10^-3^,7.45 × 10^-4^]).

**Figure 3:**
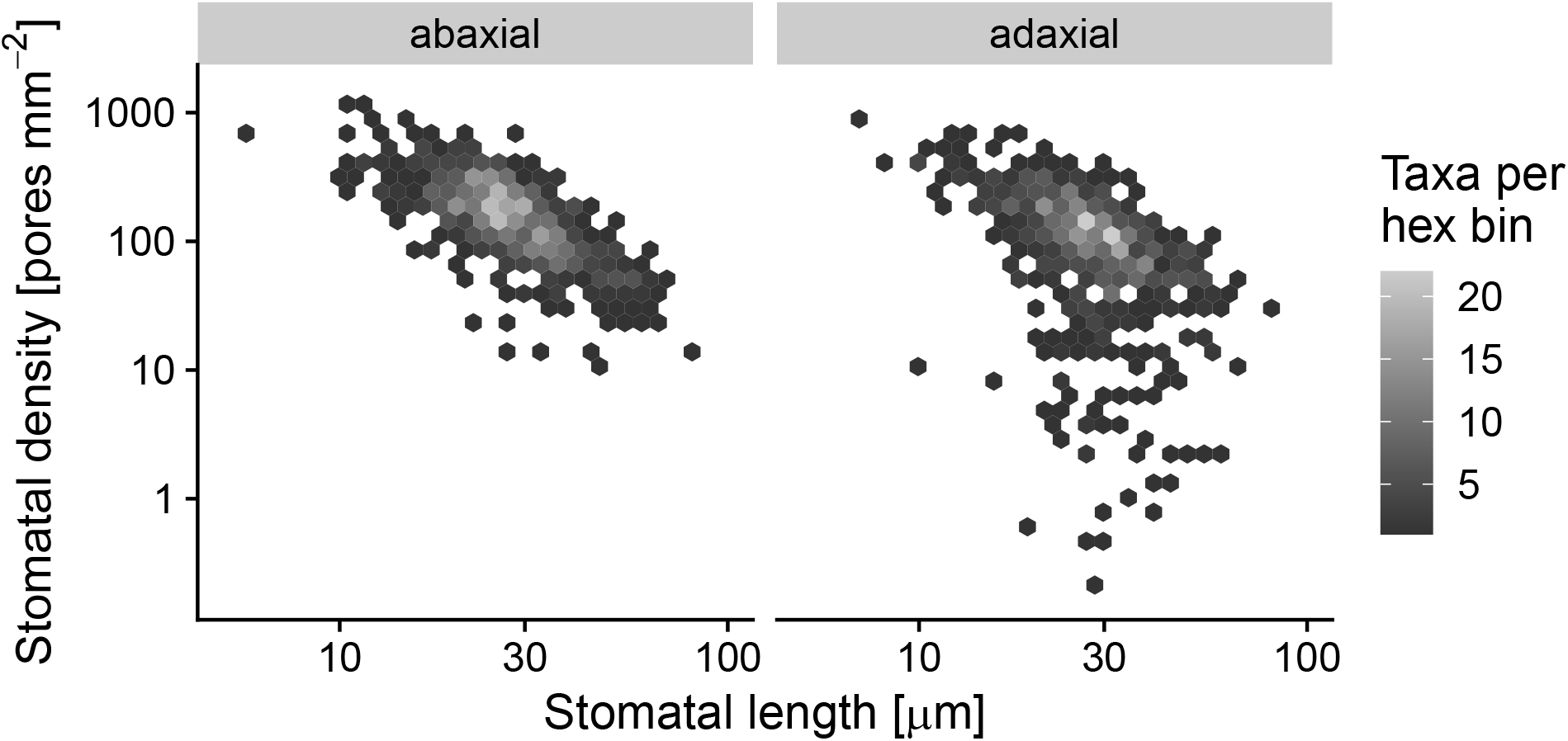
Inverse size-density scaling differs between the surfaces of amphistomatous leaf, indicating different phenotypic constraints. The panels show the relationship between stomatal length (x-axis) and stomatal density (y-axis) on a log-log scale for values measured on the abaxial leaf surface (left) and the adaxial leaf surface (right) across 638 taxa.

**Figure 4:**
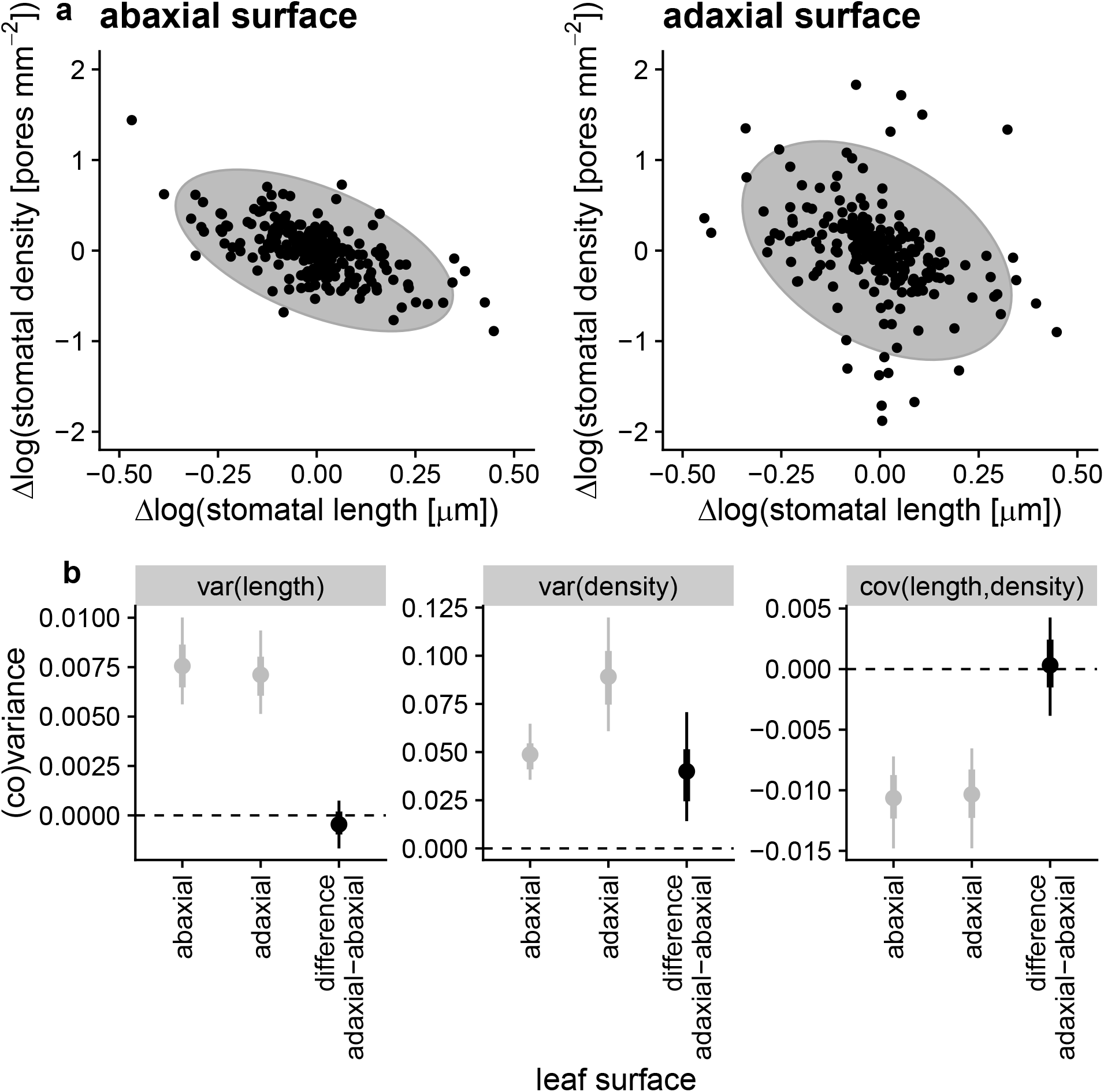
Greater overall phenotypic constraint on abaxial stomatal anatomy is evinced by more variable evolutionary divergence in adaxial stomatal density, but covariance between density and length is similar on both surfaces. (a) Data from 236 phylogenetically independent contrasts of change in log(stomatal length) (*x*-axis) and log(stomatal density) (*y*-axis) for abaxial (left panel) and adaxial (right panel) leaf surfaces. Each contrast is shown by black points and every contrast appears on both panels. Grey ellipses are the model-estimated 95% covariance ellipses. The negative covariance is similar for both surfaces but the breadth in the *y*-direction is larger for adaxial traits, indicating greater evolutionary divergence in log(stomatal density). (b) Parameter estimates (points), 66% (thick lines), and 95% HPD intervals for estimates of trait (co)variance. Grey points and lines represent ab- and adaxial values; black points and lines represent the estimated difference in (co)variance between surfaces. Only the variance for stomatal density (middle panel) is significantly greater for the adaxial surface (95% HPD interval does not overlap the dashed line at 0). Reported parameter estimates are conditioned on zero time since divergence between taxa (see Results).

### Genome size is associated with stomatal length on both surfaces

We analyzed a smaller set of 79 contrasts with data on both genome size and stomatal anatomy. Consistent with previous studies (Beaulieu et al. 2008; Jordan et al. 2015; Simonin and Roddy 2018), increased genome size was associated with increased stomatal length on both surfaces (Fig. A2). The association between genome size and stomatal density was negative, as expected, but weaker. Only the slope for adaxial stomatal density was significantly less than 0 (Fig. A2).

### Stomatal density on each surface is less integrated than stomatal length

The relationship between stomatal density on each leaf surface is visually more variable than that for stomatal length (Fig. 5). This pattern occurs because the slope and strength of integration for stomatal density on each surface is much weaker than that for stomatal length. The SMA slope between Δlog(*D*_ad_) and Δlog(*D*_ab_) is less than 1 (estimated slope = 0.742, 95% HPD interval [0.619,0.883]) and the strength of association is weakly positive (estimated *r*^2^ = 0.113, 95% HPD interval [0.0431,0.205]; Fig. 6). In contrast, the relationship between Δlog(*L*_ad_) and Δlog(*L*_ab_) is isometric (estimated slope = 1.03, 95% HPD interval [0.955,1.12]) and strongly positive (estimated r^2^ = 0.762, 95% HPD interval [0.691,0.82]; Fig. 6).

**Figure 5:**
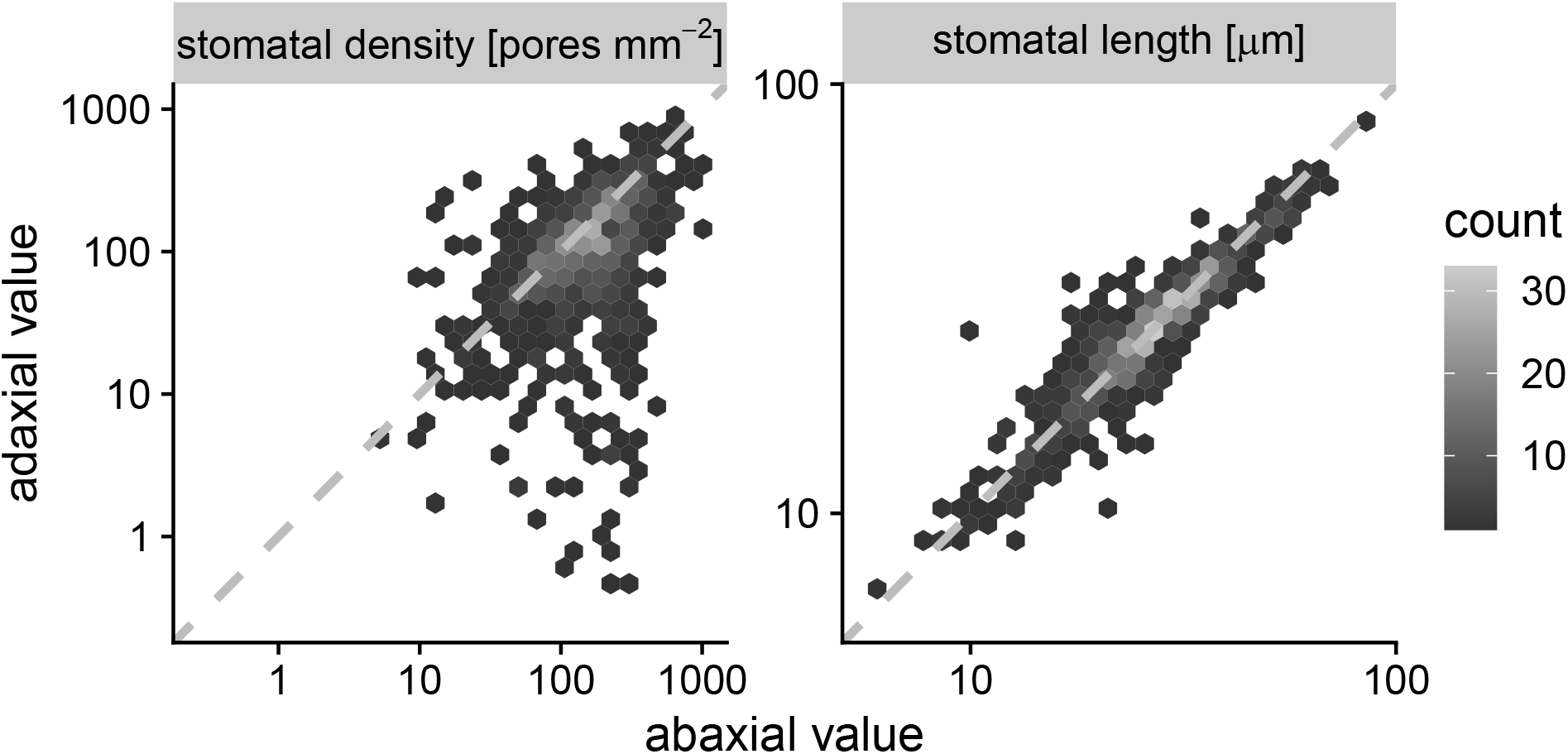
Stomatal density on each surface is less constrained (lower correlation) than stomatal length (higher correlation). Relationship between stomatal density and length on each leaf surface in a synthesis of amphistomatous leaf traits across 638 taxa. The panels show the relationship between the abaxial trait value (*x*-axis) and the adaxial trait value (*y*-axis) on a log-log scale for stomatal density (left) and stomatal length (right). The dashed line in across the middle is the 1:1 line for reference.

**Figure 6:**
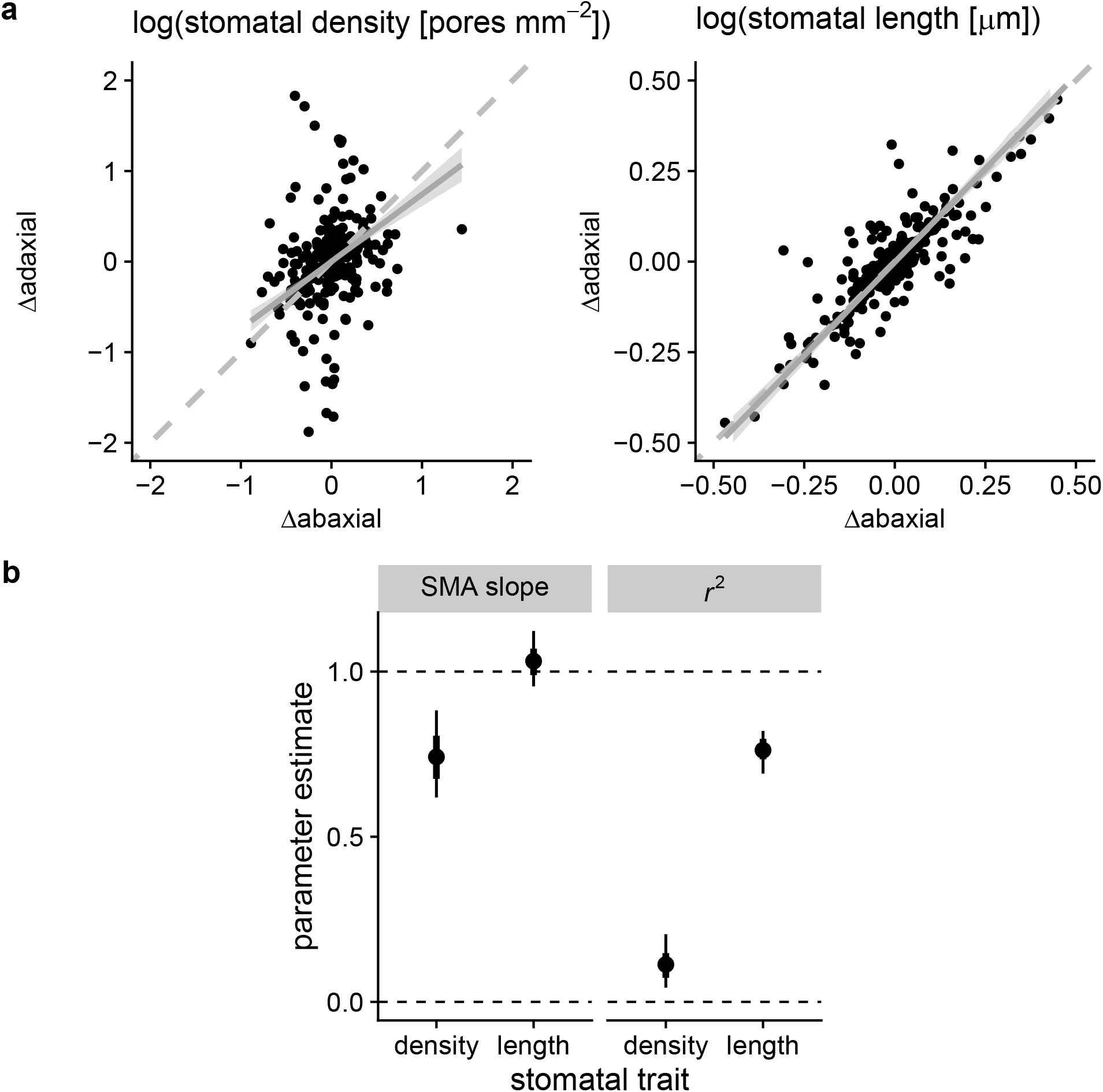
Developmental integration in stomatal length is much stronger than stomatal density between the surfaces of amphistomatous leaves (a) Data from 236 phylogenetically independent contrasts of change in the abaxial trait value (x-axis) against change in the adaxial trait value (*y*-axis) for log(stomatal density) (left panel) and log(stomatal length) (right panel). Each contrast is shown by black points and every contrast appears on both panels. Dashed grey lines are 1:1 lines for reference. Solid grey lines and ribbon the fitted SMA slope and 95% HPD interval. (b) The SMA slope (left panel) is significantly less than 1 (isometry, top dashed line) for density but very close to isometric for length. The coefficient of determination (*r*^2^, right panel) is also much greater for length than density. The points are parameter estimates with 66% (thick lines) and 95% HPD intervals. Reported parameter estimates are conditioned on zero time since divergence between taxa (see Results).

## Discussion

Functional and developmental constraints may hinder adaptation by preventing traits from evolving independently towards a multivariate phenotypic optimum. An alternative view is that phenotypic evolution is limited primarily because most phenotypes are maladaptive and selected against, which is called selective constraint. In practice, distinguishing between functional, developmental, and selective constraints is challenging. In this study, we took advantage of the fact that amphistomatous leaves produce stomata on both abaxial (usually lower) and adaxial (usually upper) surfaces. Packing limits, a functional constraint, and developmental integration should result in similar (co)variance in divergence of stomatal traits on each surface (Notes A1), whereas differing selective constraints for each surface would lead to different patterns of divergence. Although some patterns of trait (co)variance are compatible with packing limits or developmental integration (see below), independent evolution of stomatal density on each surface across flowering plants is notably inconsistent with these constraints. We therefore conclude that selective constraints, which require an adaptive explanation, most likely play a major role in the evolution of stomata and possibly other ecologically important traits.

Neither packing limits nor developmental integration were sufficient to explain two major patterns of divergence in stomatal traits (Fig. 1). Although evolutionary divergence slowed as the allocation to stomata *f_S_* increased, the effect of *f_S_* was different on each surface (Fig. 2 and A1; Table A2). The contrasting pattern of divergence on each surface is inconsistent with a common packing limit and suggests instead that selective constraints act differently on lower and upper stomata. Contrary to the developmental integration hypotheses, the greater variance in stomatal density compared to length on the adaxial surface indicates that density is more labile on this surface, though traits on each surface are not completely decoupled (Fig. 4, 6; Table A3). Consistent with the developmental integration hypotheses, divergence in stomatal length on each surface evolves isometrically at the same rate, suggesting that guard cell dimensions may not be able to evolve independently on each surface (Fig. 6). The evolutionary lability of stomatal density, despite constraints on size, show that inverse size-density scaling and bimodal stomatal ratio cannot be attributed entirely to developmental integration. Combinations of small stomata and low density that are not found on the abaxial surface are found on the adaxial surface, indicating that these rare trait combinations are developmentally accessible.

While phylogenetic comparisons usually cannot prove that phenotypic variation is adaptive, ruling out alternative hypotheses as the sole explanation is an important step toward quantifying the relative importance of selective constraint in macroevolution (McGhee 1999; Olson 2019). Establishing that traits can evolve quasi-independently is necessary but not sufficient to show that selective constraints are the primary process shaping phenotypic evolution. Packing limits and developmental integration may bias phenotypic evolution, even if they do not preclude certain stomatal trait combinations. Therefore, future research will need to combine stomatal developmental (dis)integration with biophysical models of how stomatal anatomy would vary adaptively (Olson and Arroyo-Santos 2015). Although these questions and approaches apply to any phenotype, stomata will be a useful trait because of their ecological significance and broad application to most land plants.

### Do packing limits and developmental integration lead to inverse size-density scaling?

Stomata cannot occupy more than the entire leaf surface, but realistically there is probably an upper packing limit below this hard bound. If this packing limit drives inverse size-density scaling, we should observe that divergence in stomatal size and density decrease as this limit is approached. Near the limit, large changes that reduce *f_S_* are possible, but changes that increase *f_S_* must be small so as not to exceed the limit. Furthermore, the same packing limit should apply to both ab- and adaxial leaf surfaces. Although we observe that divergence decreases with *f_S_*, the relationship is not the same on both surfaces (Fig. 2 and A1; Table A2). This implies that other factors constrain stomatal size and density before they approach a packing limit.

If meristematic cell volume and expansion integrate stomatal size and density (Brodribb, Jordan, and Carpenter 2013), then we predicted inverse size-density scaling would evolve with the same (co)variance for both ab- and adaxial leaf surfaces (Notes A1). Contrary to this prediction, there are many combinations of stomatal density and length found on adaxial leaf surfaces that are absent from abaxial leaf surfaces (Fig. 3). In principle, the different relationship between traits on each surface could be caused by different evolutionary variance in stomatal density (Var[Δlog(*D*_ab_)] ≠ Var[Δlog(*D*_ad_)]) and/or covariance (Cov[Δlog(*L*_ab_),Δlog(*D*_ab_)] ≠ Cov[Δlog(*L*_ad_),Δlog(*D*_ad_)]) on each surface. However, the covariance relationship between density and length is similar on each surface, whereas the evolutionary variance in adaxial stomatal density is significantly higher than that for abaxial density (Var[Δlog(*D*_ab_)] < Var[Δlog(*D*_ad_)]; Fig. 4). Given that the average stomatal length is usually about the same on each surface (see below), these results imply that plants can often evolve stomatal densities on each surface without a concomitant change in size. Based on our theoretical analysis, we interpret these results to mean that cell divisions affecting stomatal index are less evolutionarily constrained than the asymmetric cell division preceding the guard cell meristemoid (Fig. A3; Table A4)

The disintegration of stomatal size and density on adaxial leaf surfaces implies that the inverse size-density scaling on abaxial surfaces (Weiss 1865; Franks and Beerling 2009; de Boer et al. 2016; Sack and Buckley 2016; Liu et al. 2021) is not a developmental *fait accompli*. The lability of *D*_ad_ may explain why there is so much putatively adaptive variation in the trait along light gradients (Muir 2018) and in coordination with other anatomical traits that vary among precipitation habitats (Pathare, Koteyeva, and Cousins 2020). There may appear to be a tension between our results and recent findings that genome size, which is strongly correlated with meristematic cell volume (Šímová and Herben 2012), correlates strongly with mature guard cell size as well as the size and packing density of mesophyll cells (Roddy et al. 2020; Théroux-Rancourt et al. 2021). However, most plant species are far from their minimum cell size as determined by genome size [Roddy et al. (2020); Fig. 3]. Genome size explains 31-54% of the variation in stomatal density across the major groups of terrestrial plants (Simonin and Roddy 2018) but there is huge variation in stomatal density and stomatal length in angiosperms with rather similar genome size (*c.f*. Fig. 2 in Simonin and Roddy 2018). Genome size, a proxy for meristematic cell volume, is more strongly related to stomatal size than density (Fig. A2). Yet the decoupling of size and density on the adaxial surface suggests that meristematic cell volume is probably not a strong constraint on the final size of epidermal pavement and guard cells because of different division and expansion rates after the asymmetric cell division stage. A possible resolution is that meristematic cell volume limits the range of variation in species with exceptionally large genome, but most species can modify stomatal size and density independently of each other to optimize photosynthesis (Jordan et al. 2015; Simonin and Roddy 2018; Roddy et al. 2020; Théroux-Rancourt et al. 2021).

### Does developmental integration lead to bimodal stomatal ratio?

We predicted that if abaxial and adaxial stomata are developmentally integrated then we should observe a strong, isometric relationship between trait divergence on each surface. Consistent with this prediction, divergence in stomatal length on each surface is isometric (SMA slope = 1.03) and strongly associated (*r*^2^ = 0.762; Fig. 6). In contrast, divergence in stomatal density on each surface was not isometric (SMA slope = 0.742) and much less integrated (*r*^2^ = 0.113; Fig. 6). Since average stomatal density on each surface can evolve quasi-independently, a wide variety of stomatal ratios are developmentally possible. The stomatal developmental function is not constrained to be identical on each surface. If natural selection favored an intermediate stomatal ratio between modes, developmental processes should not constrain the expression of genetic variation to achieve it. This supports the hypothesis that the bimodal stomatal ratio pattern (Muir 2015) arises because natural selection rarely favors intermediate trait values, not because intermediate values are developmentally inaccessible.

### Limitations and future research

The ability of adaxial stomatal density to evolve independently of stomatal size and abaxial stomatal density is not consistent with packing limits or developmental integration as the primary cause leading to inverse size-density scaling or bimodal stomatal ratio. However, there are two major limitations of this study that should be addressed in future work. First, while *D*_ab_ can diverge independently of other stomatal traits globally, we cannot rule out that developmental integration is important in some lineages. Developmental constraints are often localized to particular clades but not universal (Maynard Smith et al. 1985). For example, Berg’s rule observes that vegetative and floral traits are often developmentally integrated, but integration can be broken when selection favors flowers for specialized pollination (Berg 1959, 1960; Conner and Lande 2014). Other traits evince developmental modularity, such as the independent evolution leaf and petal venation (Roddy et al. 2013). Analogously, developmental integration between stomatal anatomical traits could evolve in some lineages, due to selection or other evolutionary forces, but become less integrated in other lineages. For example, *D*_ab_ and *D*_ad_ are positively genetically correlated in *Oryza* (Ishimaru et al. 2001; Rae et al. 2006), suggesting developmental integration may contribute to low variation in stomatal ratio between species of this genus (Giuliani et al. 2013). A second major limitation is that covariation in traits like stomatal length, which appear to be developmentally integrated on each surface, could be caused by other processes. For example, since stomatal size affects the speed and mechanics of stomatal closure (Drake, Froend, and Franks 2013; Harrison et al. 2020; but see Roddy et al. 2020), there may be strong selection for similar stomatal size throughout the leaf to harmonize rates of stomatal closure. Coordination between epidermal and mesophyll development may also constrain how independently stomatal traits on each surface can evolve (Dow, Berry, and Bergmann 2017; Lundgren et al. 2019; Théroux-Rancourt et al. 2021; Borsuk et al. 2022).

Future research should identify the mechanistic basis of developmental disintegration between *D*_ab_ and *D*_ad_. Multiple reviews of stomatal development conclude that stomatal traits are independently controlled on each surface (Lake, Woodward, and Quick 2002; Bergmann and Sack 2007), but we do not know much about linkage between ab-adaxial polarity and stomatal development (Kidner and Timmermans 2010; Pillitteri and Torii 2012). Systems that have natural variation in stomatal ratio should allow us to study how developmental disintegration evolves. Quantitative genetic studies in *Brassica oleracea* L., *Oryza sativa* L., *Populus trichocarpa* Torr. & A. Gray ex Hook., *Populus* interspecific crosses, and *Solanum* interspecific crosses, typically find partial independence of *D*_ab_ and *D*_ad_; some loci affect both traits, but some loci only affect density on one surface and/or genetic correlations are weak (Ishimaru et al. 2001; Ferris et al. 2002; Hall et al. 2005; Rae et al. 2006; Laza et al. 2010; Chitwood et al. 2013; McKown et al. 2014; Muir, Pease, and Moyle 2014; Porth et al. 2015; Fetter, Nelson, and Keller 2021). For example, *Populus trichocarpa* populations have putatively adaptive genetic variation in *D*_ad_. Populations are more amphistomatous at Northern latitudes with shorter growing seasons that may select for faster carbon assimilation (McKown et al. 2014; Kaluthota et al. 2015; Porth et al. 2015). Genetic variation in key stomatal development transcription factors is associated with latitudinal variation in *D*_ad_, which should help reveal mechanistic basis of developmental disintegration between surfaces (McKown et al. 2019).

## Data avaibility

The final data set and phylogeny used in the analysis are included in the Online Appendix. The raw anatomical data and source code are deposited on Dryad (https://datadryad.org/stash/share/Tk-7ZGwhdxyxAJu2dTsBdWDaId9qmog2fGYLWgGzPwU.)

## Online Appendix

**Figure A1:**
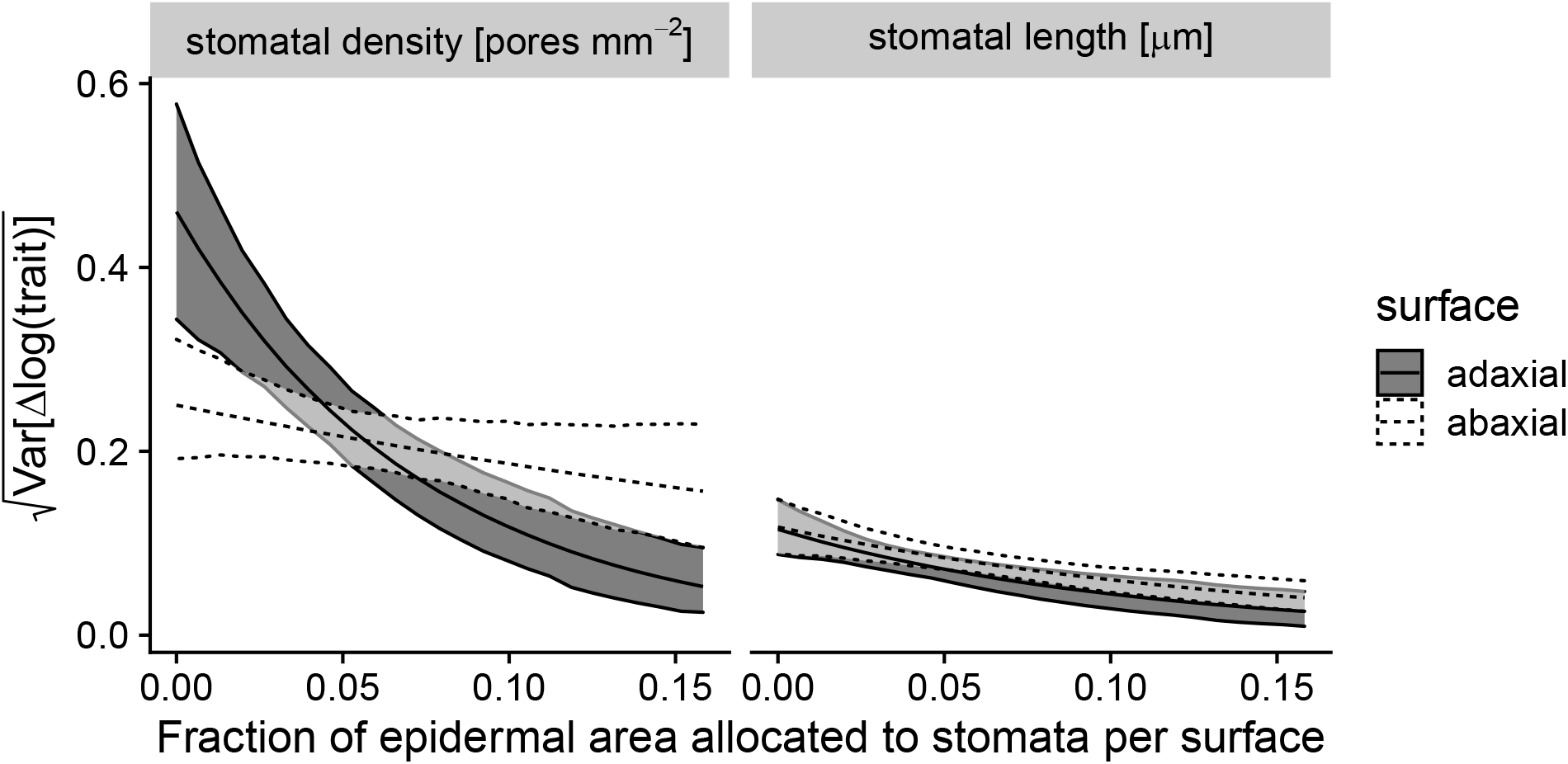
The variance in evolutionary divergence (*y*-axis, Var[Δlog(trait)]) declines the fraction of epidermal space allocated to stomata per surface (*x*-axis, *f_S_*) increases. Within each ribbon, the middle line is the median estimate and the outer lines are the 95% HPD intervals. The slope is significantly less than 0 for adaxial stomatal density and stomatal length of both surface (Table A2). Results for the standard deviation, which is the square-root of the variance, are shown.

**Figure A2:**
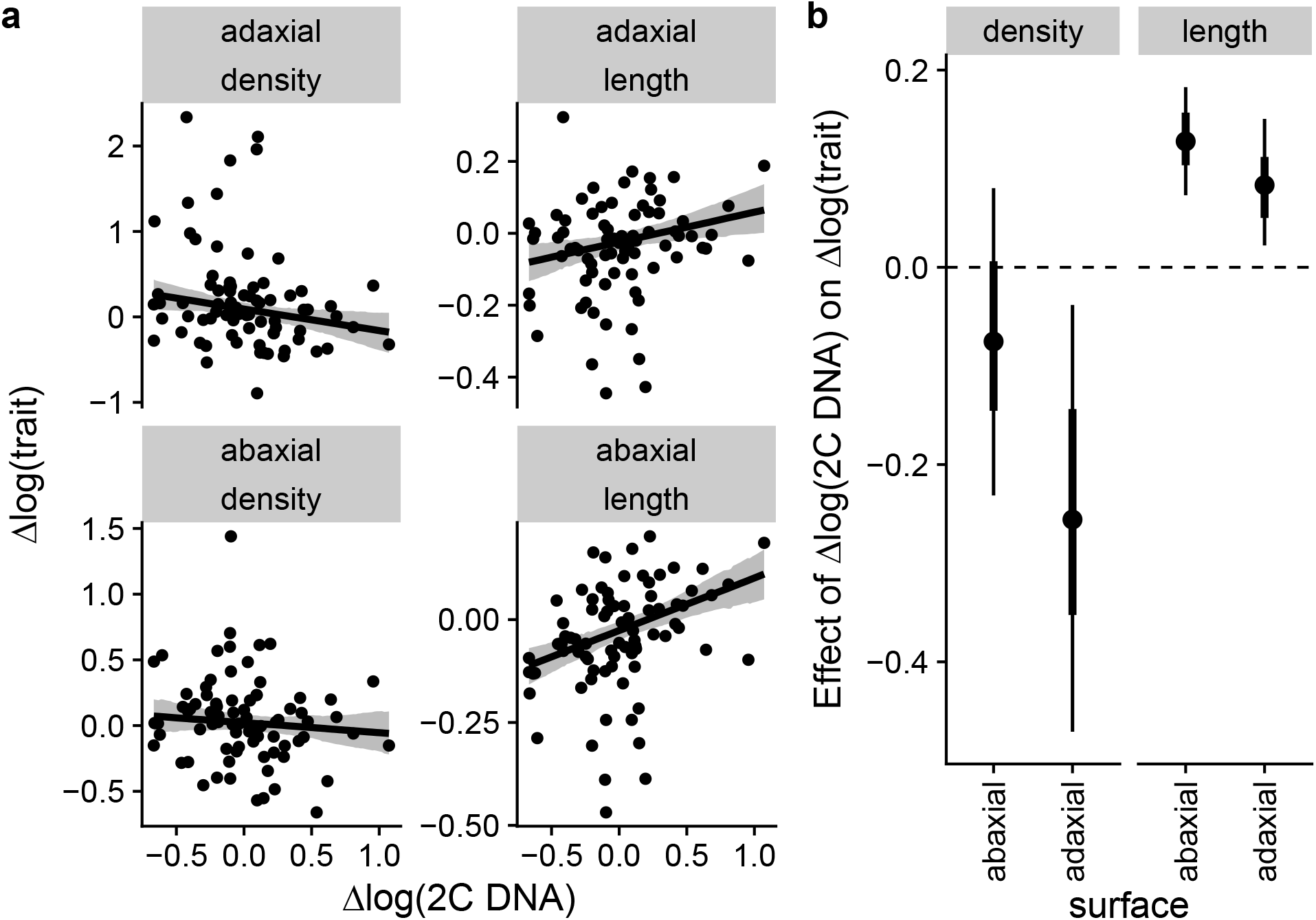
Evolutionary divergence in genome size (2C DNA content) is associated with guard cell length, but less so stomatal density. (a) Data from 79 phylogenetically independent contrasts of change in log(2C DNA) (*x*-axis) and change in log(trait) (*y*-axis) for abaxial (lower panels) and adaxial (upper panels) leaf surfaces. Each contrast is shown by black points and every contrast appears on all panels. Black lines are the median predicted trait divergence and grey ribbons are the model-estimated 95% HPD confidence bands. (b) Parameter estimates (points), 66% (thick lines), and 95% HPD intervals for estimates of the effect of change in log(2C DNA) on change in log(trait). HPD intervals that do not overlap zero indicate that divergence in genome size is associated with divergence in stomatal anatomy. Reported parameter estimates are conditioned on zero time since divergence between taxa (see Results).

Table A1: Final data set of 236 taxon pairs for analysis. `tree_node` is the node of the common ancestor of the taxon pair `sp1` and `sp2` in the phylogeny (Notes A3). `pair_age` is the time in millions of years since taxa split. The remaining columns are the trait divergence (log-scale) between taxa (Δlog(trait)).

Table A1 is a csv file uploaded with this manuscript and will be included as an online supplement upon publication.

**Table A2:** Parameter estimates and 95% highest posterior density (HPD) intervals for the effect of *f_S_* on trait divergence. For each trait (*D*_ab_, *D*_ad_, *L*_ab_, *L*_ad_) we estimated the coefficient of *f_S_* on the standard deviation of Δlog(trait) on a log-link scale. Other model parameter estimates and confidence intervals can be found in the saved model output located in the archived online repository (see Data avaibility).

| Trait(s) | Estimate | 95% HPD interval |
| --- | --- | --- |
| Effect of $f_S$ on standard deviation of $\Delta\log(\text{trait})$<br>log-link scale | | |
| $D_{ab}$ | -3.2 | $[-6.6, 0.75]$ |
| $D_{ad}$ | -12.0 | $[-17, -6.9]$ |
| $L_{ab}$ | -6.7 | $[-10, -2.9]$ |
| $L_{ad}$ | -9.8 | $[-16, -4.1]$ |

**Table A3:** Parameter estimates and 95% highest posterior density (HPD) intervals for the (co)variance of trait divergence. For each trait (*D*_ab_, *D*_ad_, *L*_ab_, *L*_ad_) we estimated the average (median) divergence between taxon pairs, denoted Δlog(trait). See Table 1 for symbol definitions. The second section is the standard deviation of Δlog(trait). The third section is the estimated coefficient of pair age (millions of years) on the standard deviation on a log-link scale. The fourth section is the estimated correlation coefficient between Δlog(trait) of all pairwise trait combinations. The final section is the estimated *v* family of the Student *t* distribution.

| Trait(s) | Estimate | 95% HPD interval |
| --- | --- | --- |
| Average $\Delta\log(\text{trait})$ | | |
| $D_{ab}$ | 0.00180 | $[-0.029, 0.035]$ |
| $D_{ad}$ | 0.00069 | $[-0.045, 0.047]$ |
| $L_{ab}$ | -0.00620 | $[-0.019, 0.0079]$ |
| $L_{ad}$ | -0.00760 | $[-0.021, 0.0055]$ |
| Standard deviation of $\Delta\log(\text{trait})$ | | |
| $D_{ab}$ | 0.220 | $[0.19, 0.26]$ |
| $D_{ad}$ | 0.300 | $[0.25, 0.35]$ |
| $L_{ab}$ | 0.087 | $[0.075, 0.1]$ |
| $L_{ad}$ | 0.084 | $[0.073, 0.098]$ |
| Effect of pair age on standard deviation of $\Delta\log(\text{trait})$<br>log-link scale | | |
| $D_{ab}$ | 0.004 | $[-0.0034, 0.013]$ |
| $D_{ad}$ | 0.013 | $[0.0047, 0.022]$ |
| $L_{ab}$ | 0.012 | $[0.0057, 0.02]$ |
| $L_{ad}$ | 0.013 | $[0.0064, 0.019]$ |
| Correlation between $\Delta\log(\text{trait})$ | | |
| $D_{ab} - D_{ad}$ | 0.34 | $[0.21, 0.45]$ |
| $D_{ab} - L_{ab}$ | -0.56 | $[-0.65, -0.45]$ |
| $D_{ab} - L_{ad}$ | -0.46 | $[-0.57, -0.35]$ |
| $D_{ad} - L_{ab}$ | -0.37 | $[-0.48, -0.25]$ |
| $D_{ad} - L_{ad}$ | -0.41 | $[-0.52, -0.31]$ |
| $L_{ab} - L_{ad}$ | 0.87 | $[0.83, 0.91]$ |
| Student $t$ family parameter $\nu$ | | |
| — | 3.4 | $[2.5, 4.4]$ |

### Notes A1: Theory connecting developmental function, constraint, and integration

Below we provide a conceptual background to motivate the derivation of a stomatal developmental function. We then derive predictions for how stomatal size and density should diverge with or without developmental constraint. We then explain why comparing evolutionary divergence of lower and upper stomatal anatomy provides an important additional line of evidence on the contribution of developmental integration to phenotypic macroevolution. Fig. A3 is a graphical summary of our analysis.

#### Conceptual background

Developmental integration in stomatal anatomy is plausible because epidermal pavement cells and stomata share an early developmental history, originating from the same leaf meristem tissue. If all other factors are held constant, meristematic cell volume, which is largely determined by genome size (Šímová and Herben 2012), and early expansion rate increase both epidermal cell and stomatal area proportionally. This mechanically decreases stomatal density because the same number of stomata per epidermal cell (stomatal index) are spread farther apart by larger epidermal cells. Developmental integration between stomatal size and density arises naturally if meristematic cell volume and/or expansion rate evolve, but the remaining steps of stomatal development are fixed. As described in detail below, we mathematically formalize these later steps in stomatal development into a ‘developmental function’ inspired by Wagner (1989). Wagner’s used a developmental function to map genetic variance onto phenotypic variance. The developmental function can cause a disposition for phenotypic covariance, depending on the amount of pleiotropy. For example, genetic changes in a growth factor could be highly pleiotropic, simulta-neously altering the size of many tissues. Wagner used the developmental function to model microevolution, but if we suppose that the developmental function is fixed over long time periods, it can be used to predict macroevolutionary divergence under developmental constraint. If the developmental function is fixed or highly constrained, species may never possess the genetic variation to access regions of phenotypic space. If the developmental function itself can evolve readily, then traits should be able to evolve independently given sufficient time for mutation, selection, and divergence. Finding that the developmental function is malleable would lend less credence to the importance of developmental constraint and lend more credence to selective hypotheses.

The stomatal developmental function is probably not fixed, potentially allowing for independent evolution of stomatal size and density. The conceptual model of stomatal development by Dow and Bergmann (2014) identifies three key cell division types that could shape stomatal density and size. First, asymmetric division of undifferentiated epidermal cells forms the guard cell meristemoid. Larger allocation to and/or greater expansion of the meristemoid as it matures to a guard mother cell increases stomatal size without affecting density. Second, spacing divisions in developing epidermal cells increase stomatal density and index while maintaining spacing. Third, amplifying divisions generate more epidermal cells without further differentiation of stomata, decreasing stomatal density and index. Changing the probability of spacing and amplifying divisions affects stomatal density without changing size.

Below we formalize these models of developmental (dis)integration to address the following two questions:

1. How would stomatal size and density (co)diverge if the developmental function is fixed? We refer to this as the ‘developmental integration’ hypothesis.
2. How would stomatal size and density (co)diverge if the developmental function is not fixed? We refer to this as the ‘developmental disintegration’ hypothesis.

#### Theory

##### A developmental function for stomatal size and density

In this section we derive a stomatal developmental function by extending the model of Sack and Buckley (2016) in two ways. First, we provide an explicit, albeit simple, map from meristematic cell volume to stomatal size and density. Second, we use random variable algebra (Lynch and Walsh 1998) to derive expectations for the variance in stomatal anatomy among species. Sack and Buckley (2016) consider three anatomical properties of a leaf surface, the projected epidermal cell area E, the area of the stomatal apparatus S, and the stomatal index *I*:

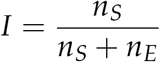

*n_S_* and *n_E_* are the number of stomatal and other epidermal cells, respectively, on the leaf surface. Throughout this we appendix we focus on stomatal size (*S*) rather than guard cell length (*L*) because it is mathematically simpler. For comparison with our data on *L*, we derive predictions using the fact that *S* = *jL*^2^ where *j* = 0.5 for non-grasses and 0.125 for grasses (Sack and Buckley 2016).

Next, we assume that the area of epidermal cells and stomata are proportional to the meristematic cell volume *M*:

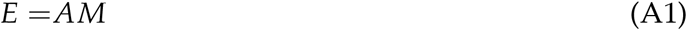

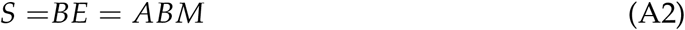

The coefficient *A* is determined by the early cell expansion and division rates, which we do not model explicitly. *B* is determined by the placement of the asymmetric cell division generating the guard mother cell (Bergmann and Sack 2007) and subsequent expansion of the guard cell meristemoid. For example, in *Arabidopsis thaliana*, the cell volume of shoot meristematic cells is approximately 200 *μ*m^3^ (Price, Sparrow, and Nauman 1973) and the epidermal and stomatal sizes are roughly 1000 and 250 *μ*m^2^ (Dow, Bergmann, and Berry 2014). Therefore 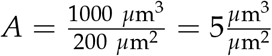 and 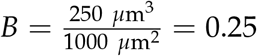.

Following Sack and Buckley (2016) the stomatal density as a function of *E, S*, and *I* is:

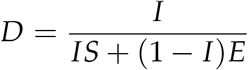

For analytical tractability, we use he first-order Taylor series approximation around *I* = 0 because *I* is typically much closer to 0 than 1:

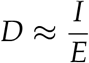

Below we show that this approximation accurately models the correlation in divergence between stomatal size and density by comparing it to random simulations (Fig. @ref{fig:check-approximation}.

Substituting Eqn. A2 into the above expression we obtain

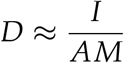

Now we can can derive a developmental function to map from *M* to *S* and *D*. We assume that *M* is determined by genome size (Šímová and Herben 2012) and, possibly, other genetic and environmental factors that we do not track explicitly in our model. As with our empirical analysis, we work with the log-transformed values of *S* and *D* to linearize the developmental function. For brevity, let the lowercase variables be the log-transformed values of their uppercase counterparts (e.g. *d* = log(*D*)). With these assumptions, we obtain:

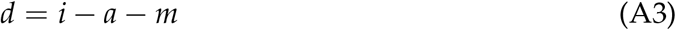

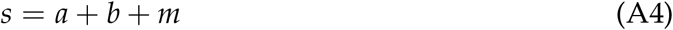

#### Hypotheses

To address the two overarching questions posed above, we will use the theory in the previous section to derive predictions for two hypotheses. The developmental integration hypothesis can be thought of as a null hypothesis for how stomatal size and density diverge when the developmental function is fixed. The second hypothesis relaxes this constraint.

1. Developmental integration hypothesis: the stomatal developmental function is fixed; divergence in stomatal size and density is caused only by divergence in meristematic cell volume and early expansion rate.
2. Developmental disintegration hypothesis: the stomatal developmental function is not fixed; divergence in stomatal size and density is caused by the combined divergence in meristematic cell volume, early expansion, and later cell divisions, the asymmetric, spacing, amplifying divisions discussed in the Conceptual background.

##### Developmental integration hypothesis

Under this hypothesis, meristematic cell volume and expansion rate integrate stomatal size and density because the developmental function is constrained. We know that meristematic cell volume can evolve as a product of genome size, so a natural null hypothesis is that *m* varies but the developmental parameters *a*, *b*, and *i* in Eqn. A4 are constant or vary little relative to *m*. Let the divergence between taxa *i* and *j* be:

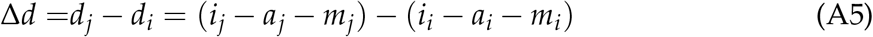

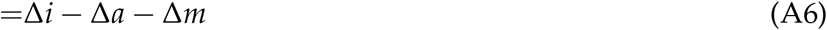

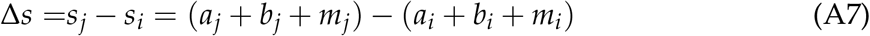

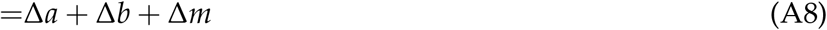

When developmental parameters are fixed Δ*a* = Δ*b* = Δ*i* = 0. This leads to integration between *s* and *d* mediated by *m* because Δ*s* = Δ*m*, Δ*d* = –Δ*m*, and Cov[Δ*s*, Δ*d*] = – Var[Δ*m*]. Strong developmental integration would also persist if Δ*b* = Δ*i* = 0 but Δ*a* ≠ 0. In that case, Δ*s* = Δ*a* + Δ*m*, Δ*d* = – (Δ*a* + Δ*m*), and Cov[Δ*s*, Δ*d*] = –Var[Δ*a* + Δ*m*]. In either case, the correlation between between Δ*d* and Δ*s* is –1 because –Cov[Δ*s*, Δ*d*] = Var[Δ*d*] = Var[Δ*s*]:

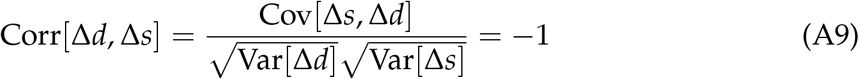

In summary, developmental constraint on stomatal index and allocation to guard mother cells during asymmetric cell division leads to developmental integration between stomatal size and density. Developmental integration can be mediated by either meristematic cell volume and/or epidermal cell expansion since they are colinear.

##### Developmental disintegration hypothesis

Here we show that developmental disintegration is mediated by divergence in stomatal index and asymmetric cell division. In conceptual models of stomatal development (Dow and Bergmann 2014), asymmetric division forms the meristemoid to the guard mother cell. After asymmetric division, spacing divisions increase stomatal density and index whereas amplifying divisions decrease both quantities. Above we assumed these processes were constrained; here we relax that assumption. First, we assume that Δ*b* = 0 and Δ*i* ≠ 0. Further, we assume for simplicity that there is no covariance in divergence between *m* and *i* (Cov[Δ*i*, Δ*m*] = 0. Using random variable algebra, the (co)variance and correlation between divergence in stomatal density and size are:

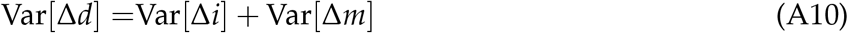

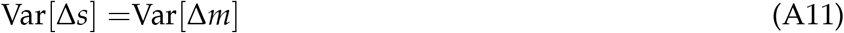

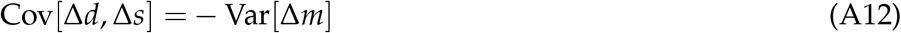

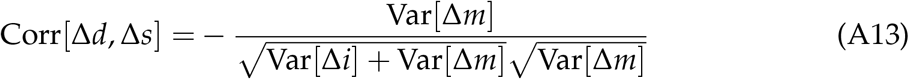

Compared to the developmental integration hypothesis, variation in stomatal index leads to greater variation in stomatal density and disintegration (lower correlation) between density and size. The approximation in Eqn. A13 matches simulated values well for realistic values of stomatal index (Fig. A4).

Next, we switch our assumptions such that Δ*b* ≠ 0 and Δ*i* = 0. We again make the simplifying assumption that there is no covariance in divergence between *m* and *b* (Cov[Δ*b*, Δ*m*] = 0. The (co)variance and correlation between stomatal density and size are:

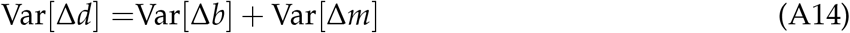

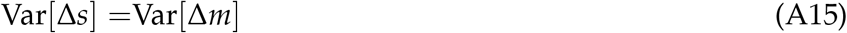

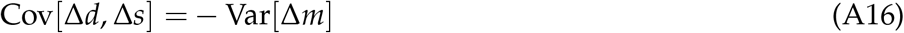

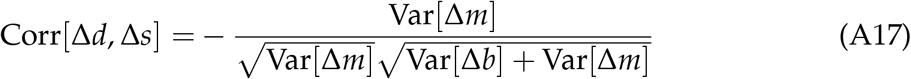

As with stomatal index, variation in asymmetric cell division also causes developmental disintegration. The key difference is that disintegration is driven by greater variation in stomatal size rather than density.

##### Predictions

In this section, we summarize the predictions for each hypothesis (Table A4) and show they can be difficult to distinguish under certain parameter combinations. Comparing the divergence of stomatal density and size on each surface provides additional evidence that can help resolve competing hypotheses. We can convert predictions from stomatal size to length using the relationship from Sack and Buckley (2016): *S* = *jL*^2^ or *s* = log(*j*) + 2*l* on the log-transformed scale. It follows that Δ*s* = 2Δ*l*, Var[Δ*s*] = 4Var[Δ*l*], and Corr[Δ*d*,Δ*s*] = Corr[Δ*d*,Δ*l*].

**Table A4:** Key predictions about the (co)variance and correlation in divergence of log(stomatal density) (Δ*d*) and log(stomatal size) (Δ*s*) under the developmental integration and disintegration hypotheses. “Single surface” predictions apply to divergence in stomatal traits on either surface; “Both surfaces” predictions compare the divergence of traits on surface to that of the other. We further contrast two variants of the disintegration hypothesis, where either stomatal index (Var[Δ*i*] ≠ 0) or asymmetric cell division (Var[Δ*b]* ≠ 0) diverges.

| Hypothesis | Predictions |  |
| --- | --- | --- |
|  | Single surface | Both surfaces |
| Developmental integration | $\text{Var}[\Delta d] = \text{Var}[\Delta s]$<br>$\text{Corr}[\Delta d, \Delta s] = -1$ | $\text{Var}[\Delta d_{ab}] = \text{Var}[\Delta d_{ad}]$<br>$\text{Var}[\Delta s_{ad}] = \text{Var}[\Delta s_{ad}]$<br>$\text{Cov}[\Delta d_{ab}, \Delta s_{ab}] = \text{Cov}[\Delta d_{ad}, \Delta s_{ad}]$ |
| Developmental disintegration<br>$\text{Var}[\Delta i] \neq 0$ | $\text{Var}[\Delta d] > \text{Var}[\Delta s]$<br>$0 < \text{Corr}[\Delta d, \Delta s] < -1$ | $\text{Var}[\Delta d_{ab}] \neq \text{Var}[\Delta d_{ad}]$<br>$\text{Var}[\Delta s_{ad}] = \text{Var}[\Delta s_{ad}]$<br>$\text{Cov}[\Delta d_{ab}, \Delta s_{ab}] \neq \text{Cov}[\Delta d_{ad}, \Delta s_{ad}]$ |
| Developmental disintegration<br>$\text{Var}[\Delta b] \neq 0$ | $\text{Var}[\Delta d] < \text{Var}[\Delta s]$<br>$0 < \text{Corr}[\Delta d, \Delta s] < -1$ | $\text{Var}[\Delta d_{ab}] = \text{Var}[\Delta d_{ad}]$<br>$\text{Var}[\Delta s_{ad}] \neq \text{Var}[\Delta s_{ad}]$<br>$\text{Cov}[\Delta d_{ab}, \Delta s_{ab}] \neq \text{Cov}[\Delta d_{ad}, \Delta s_{ad}]$ |

The sections above clarify that it is possible to use the (co)divergence in stomatal density and size to test whether developmental integration contributes to phenotypic macroevolution. The problem is that there are parameter combinations where the (co)divergence in stomatal density and size appear consistent with strong developmental integration even when there is no constraint on the developmental function. For illustration, consider an extreme example where there is no divergence in *m* or *a* (Var[Δ*m*] = Var[Δ*a*] = 0) and the (co)variance in Δ*b* and Δ*i* are aligned such that Var[Δ*b*] = Var[Δ*i*] = —Cov[Δ*b*, Δ*i*]). This leads to the same predictions as the maximally constrained model, even though there is no constraint:

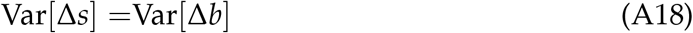

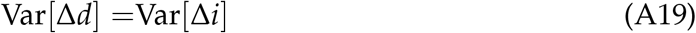

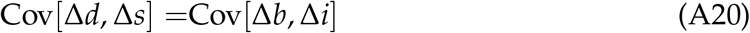

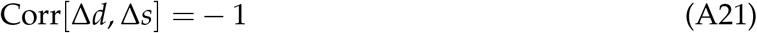

Ideally, we would measure Δ*b* and Δ*i* to test whether they contribute significantly to divergence in stomatal density and size. This is challenging because comparative data on *b* and *i* is scarcer than that for *d* and *s*. As a result, we can test whether Δ*b* and Δ*i* contribute in certain lineages but cannot directly quantify their relative importance for angiosperm macroevolution in general, as we attempt in this study.

We therefore take an alternative approach, leveraging the fact that stomatal trait evolution on each surface provides an additional line of evidence. If the stomatal developmental function is constrained then stomatal size and density on each surface should diverge in concert. Conversely, if the stomatal size and density on each surface diverge independently, this provides strong evidence that the developmental function is not fixed. If the developmental function differs between leaf surfaces with identical genomes then it seems implausible that it could not diverge over macroevolutionary time if there were selection. In that case we should give less credence to any hypothesis which posits that the developmental function *cannot* evolve.

**Figure A3:**
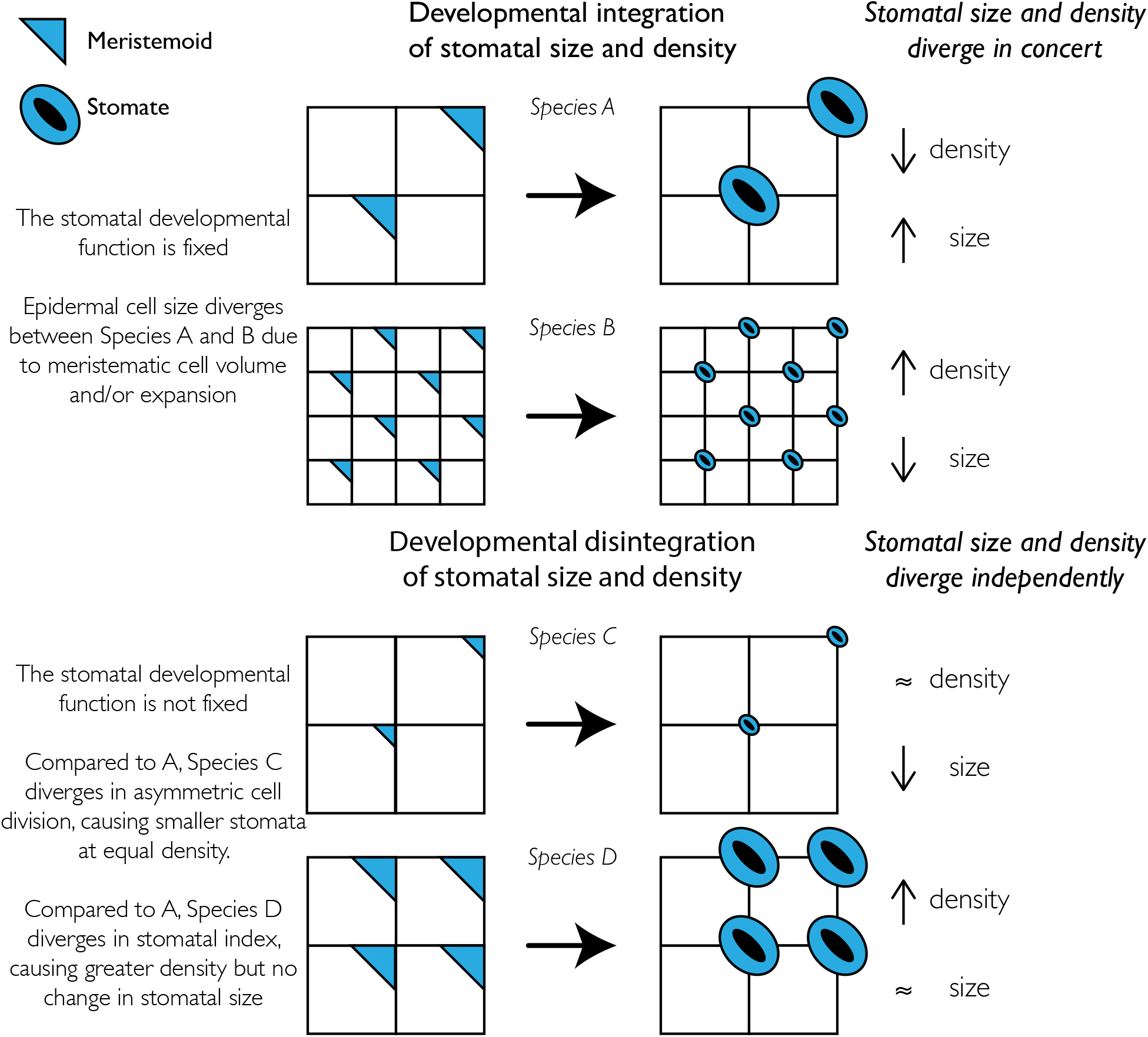
Graphical summary describing contrasting predictions of developal integration and disintegration hypotheses. Meristematic cell volume and expansion determine the epidermal (white squares) and guard meristemoid (blue triangles) cell sizes before final differentiation into stomata. Because the developmental function is fixed, larger meristematic cell volume and greater expansion result in larger stomata at lower density (Species A); smaller meristematic cell volume and less expansion result in smaller stomata at higher density (Species B). Stomatal size and density can evolve independently if the developmental function is not fixed. Species C diverges in size but not density by allocating less volume to the guard meristemoid during asymmetic cell division. Species D diverges in density but not size by increasing stomatal index.

**Figure A4:**
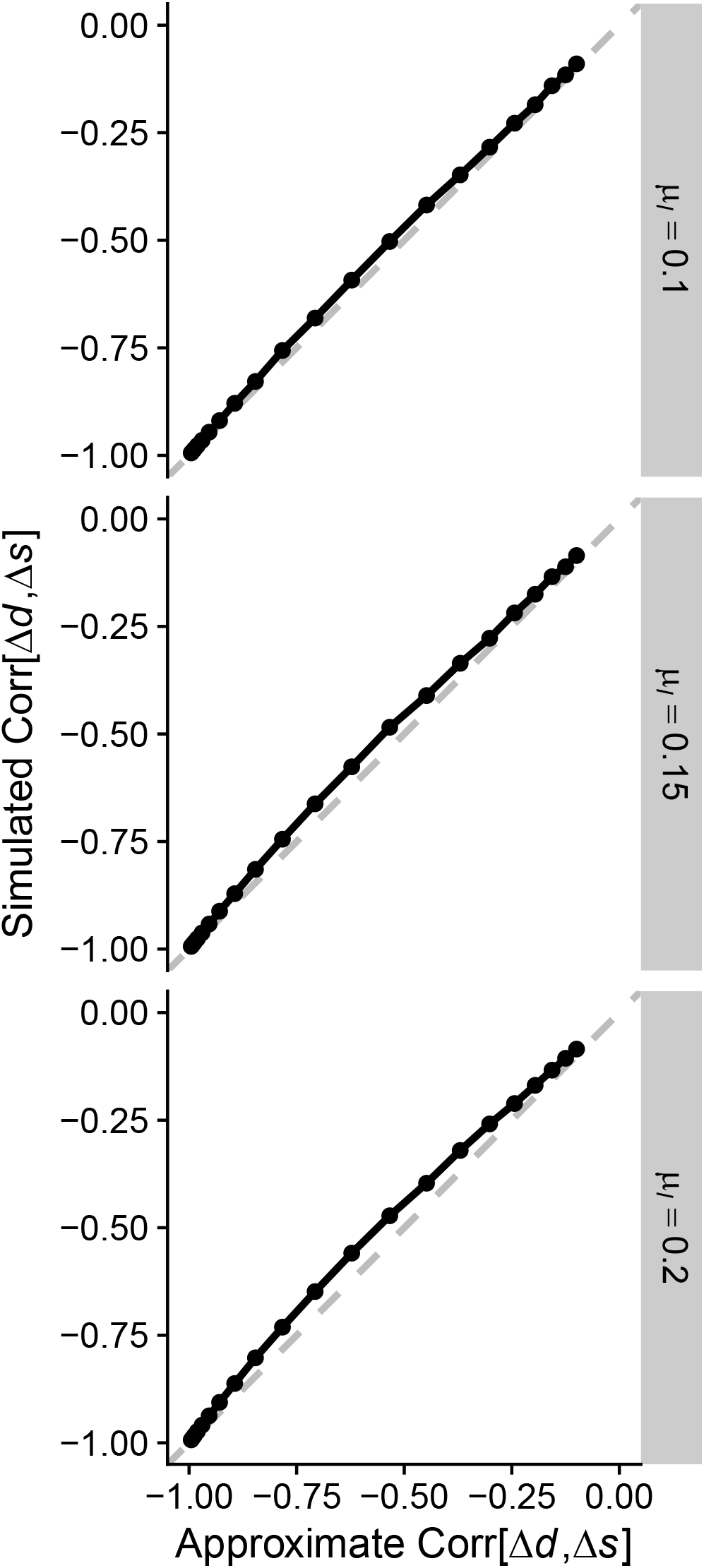
The approximation used to derive the correlation between log-transformed divergence in stomatal density and size (Corr[Δ*d*, Δ*s*] in Eqn. A13 matches simulated values. Each panel shows the relationship between approximate (*x*-axis) and calculated correlation values from 10^5^ random simulations per point (*y*-axis). The approximation is more accurate when the average stomatal index is low (*μ_I_* = 0.1) and less accurate when stomatal index is greater. Parameter values for simulations were *A* = 5, *B* = 0.25, *μ_m_* = log[200 *μ*m^3^], *σ*_Δ*i*_ = 0.1. The value of these parameters did not affect the results. The correlation changed based *σ*_Δ*m*_, which varied between 0.1 and 10 × *σ*_Δ*i*_).

### Notes A2: Phylogeny

We resolved taxonomic names using the R package **taxize** version 0.9.100 (Chamberlain and Szöcs 2013). We queried taxonomic names supplied by the original study authors on 2022-08-08 from the following sources: GRIN Taxonomy for Plants (United States Department of Agriculture, Agricultural Research Service 2020), Open Tree of Life Reference Taxonomy (Rees and Cranston 2017), The International Plant Names Index (The Royal Botanic Gardens et al. 2020), Tropicos - Missouri Botanical Garden (Missouri Botanical Garden 2020). We retained the maximum scoring matched name with taxize score ≥ 0.75 (a score of 1 is a perfect match). In 5 ambiguous cases we manually curated names. Taxonomic name resolution reduced the data set from 1120 to 1080 taxa. Most taxa are different species, but some recognized subspecies and varieties are also included. All algorithms and choices are documented in the associated source code.

We used the R packages **taxonlookup** version 1.1.5 (Pennell, FitzJohn, and Cornwell 2016) and **V.phylomaker** version 0.1.0 (Jin and Qian 2019) to maximize overlap between our data set and the GBOTB.extended mega-tree of seed plants (S. A. Smith and Brown 2018; Zanne et al. 2014). We further resolved large (≥ 4 taxa) polytomies in 29 clades with sufficient sequence data using **PyPHLAWD** version 1.0 (S. A. Smith and Walker 2019) in Python 3.9.2 (Python Software Foundation, https://www.python.org/). We used sequence data from the most recent GenBank Plant and Fungal sequences database division (Ouellette and Boguski 1997). We inferred subtree phylogenies using RAxML version 8.2.12 (Stamatakis 2014) and conducted molecular dating using the `chronos()` function in the R package **ape** version 5.6.2 (Paradis and Schliep 2019) to obtain ultrametric trees. We grafted resolved, ultrametric subtrees onto the mega-tree at the polytomy nodes and rescaled to keep the mega-tree ultrametric. In some cases, resolving polytomies was not possible because there was little or no overlap between taxa in the data set and taxa with sequence data available for **PyPHLAWD**. In these cases, we randomly selected two taxa as a phylogenetially independent pair and dropped the rest. Remaining polytomies of three taxa were resolved randomly using the `multi2di()` function in **ape**. The final data set for which we had both trait and phylogenetic information contained 638 taxa (Notes A3).

**Table A5:** Primary sources of stomatal anatomical data and the taxa covered by each source.

| Source | Taxa |
| --- | --- |
| Arambarri et al. (2005) | lotus |
| Avita and Inamdar (1980) | ranunculaceae,paeoniaceae |
| Bucher et al. (2017) | many |
| Caldera et al. (2017) | arabidopsis thaliana |
| Chandra (1967) | solanum |
| Conesa et al. (2019) | limonium |
| Eckerson (1908) | many |
| Gindel (1969) | many |
| Giuliani et al. (2013) | oryza |
| Hanafy et al. (2019) | mentha |
| Huang (2019) | trees |
| Juhász (1966) | solanum |
| Juhász (1968) | solanum |
| Kannabiran and Ramassamy (1988) | apocynaceae |
| Kawamitsu et al. (1996) | grasses |
| Khan et al. (2019) | gymnosperms |
| Kim (1987) | silverswords |
| McKown, Akamine, and Sack (2016) | scaevola |
| Muir, Galmés, and Conesa (2022) | solanum |
| Pallardy and Kozlowski (1979) | populus |
| Pandey and Nagar (2003) | many |
| Pathare, Koteyeva, and Cousins (2020) | grasses |
| Rivera, Villaseñor, and Terrazas (2017) | asteraceae |
| Rodriguez (2021) | eucalyptus |
| Scalon et al. (2016) | passovia |
| Siddiqi, Ahmad, and Rehman (1991) | euphorbiaceae |
| Sporck (2011) | euphorbia |
| Stenglein et al. (2003a) | lotus |
| Stenglein et al. (2003b) | lotus |
| Sundberg (1986) | many |
| Szymura and Wolski (2011) | solidago |
| Xiong and Flexas (2020) | many |
| Yang et al. (2014) | many |
| Zarinkamar (2006) | monocots |
| Zarinkamar (2007) | eudicots |
| Zhao et al. (2020) | monocots |
| Zlatković et al. (2017) | sedum |
| Zoric et al. (2009) | trifolium |

## Notes

https://github.com/cdmuir/stomata-independence

https://github.com/cdmuir/ropenstomata

### Competing Interest Statement

The authors have declared no competing interest.

### Summary of Updates

Abstract, Discussion, and Figure captions revised to clarify main conclusions and advance. Minor changes and other grammatical fixes.

